# Early changes in synaptic and intrinsic properties of dentate gyrus granule cells in a mouse model of Alzheimer’s disease neuropathology and atypical effects of the cholinergic antagonist atropine

**DOI:** 10.1101/2020.09.30.321646

**Authors:** David Alcantara-Gonzalez, Elissavet Chartampila, Helen E Scharfman

**Author notes:** **Corresponding author:** Helen E. Scharfman, E-Mail, The Nathan Kline Institute, Center for Dementia Research, 140 Old Orangeburg Rd. Bldg. 35, Orangeburg, NY 10962. Declarations of interest: none. **E-mail address of authors: DAG:**; **EC:**. **HES:**.

## Abstract

It has been reported that hyperexcitability occurs in a subset of patients with Alzheimer’s disease (AD) and hyperexcitability could contribute to the disease. Several studies have suggested that the hippocampal dentate gyrus (DG) may be an important area where hyperexcitability occurs. Therefore, we tested the hypothesis that the principal DG cell type, granule cells (GCs), would exhibit changes at the single-cell level which would be consistent with hyperexcitability and might help explain it. We used the Tg2576 mouse, where it has been shown that hyperexcitability is robust at 2-3 months of age. GCs from 2-3-month-old Tg2576 mice were compared to age-matched wild type (WT) mice. Effects of muscarinic cholinergic antagonism were tested because previously we found that Tg2576 mice exhibited hyperexcitability *in vivo* that was reduced by the muscarinic cholinergic antagonist atropine, counter to the dogma that in AD one needs to boost cholinergic function. The results showed that GCs from Tg2576 mice exhibited increased frequency of spontaneous excitatory postsynaptic potentials/currents (sEPSP/Cs) and reduced frequency of spontaneous inhibitory synaptic events (sIPSCs) relative to WT, increasing the excitation:inhibition (E:I) ratio. There was an inward glutamatergic current that we defined here as a novel synaptic current (nsC) in Tg2576 mice because it was very weak in WT mice. Although not usually measured, intrinsic properties were distinct in Tg2576 GCs relative to WT. In summary, GCs of the Tg2576 mouse exhibit early electrophysiological alterations that are consistent with increased synaptic excitation, reduced inhibition, and muscarinic cholinergic dysregulation. The data support previous suggestions that the DG and cholinergic system contribute to hyperexcitability early in life in AD mouse models.

**HIGHLIGHTS:**

- Granule cells (GCs) in young Tg2576 mice had abnormal synaptic activity.
- Young Tg2576 GCs had increased excitatory and reduced inhibitory synaptic activity.
- GC intrinsic properties were altered in young Tg2576 mice.
- Muscarinic cholinergic regulation of GCs was altered in young Tg2576 mice.
- Atropine led to a novel spontaneous inward current that was very strong in Tg2576 GCs.

## INTRODUCTION

Alzheimer’s disease (AD) is a neurodegenerative and progressive disorder characterized by increasing impairment in learning and memory with age, and two major types of neuropathology: amyloid-β (Aβ) plaque and neurofibrillary tangles (Braak and Braak, 1991; Lue et al., 1999; Walsh and Selkoe, 2007; Wang et al., 2013). The disease is common, has devastating effects on patients, and there is no cure. As a result, it is increasingly important to understand the contributing factors so that new treatments can be developed.

Several lines of evidence suggest that hyperexcitability is a contributing factor, primarily in familial AD (Chin and Scharfman, 2013; Friedman et al., 2012; Ghatak et al., 2019; Noebels, 2011; Nygaard et al., 2015; Palop et al., 2007; Palop and Mucke, 2009; Palop and Mucke, 2010; Petrache et al., 2019; Siskova et al., 2014; Styr and Slutsky, 2018; Verret et al., 2012; Vossel et al., 2013; Vossel et al., 2017). Hyperexcitability is manifested by intermittent seizures or epileptiform activity in an electroencephalogram (EEG) (Chin and Scharfman, 2013; Friedman et al., 2012; Gureviciene et al., 2019; Noebels, 2011; Nygaard et al., 2015; Palop et al., 2007; Palop and Mucke, 2009; Palop and Mucke, 2010; Verret et al., 2012; Vossel et al., 2013; Vossel et al., 2017). The seizures are easily missed because they are not always accompanied by movement, and epileptiform activity is easily missed because an EEG is not always conducted, or recordings are made far from the site of abnormal activity (Lam et al., 2019). For these reasons and others, there are still many questions about the extent to which hyperexcitability occurs and its importance to AD pathophysiology (Friedman et al., 2012; Leonard and McNamara, 2007; Scarmeas et al., 2009; Scharfman, 2019; Vossel et al., 2017).

Hyperexcitability also occurs in mouse models of AD (Chin and Scharfman, 2013; Palop et al., 2007; Palop and Mucke, 2009; Scharfman, 2012), which is notable because the mice primarily simulate the subset of AD that is familial, rather than all types of AD (Gotz et al., 2018; Gotz and Ittner, 2008; Scharfman, 2012). The mouse models have led to many insights into the nature of hyperexcitability and the potential to rescue cognitive impairments by reducing the heightened neuronal activity (Bakker et al., 2012; Gheyara et al., 2014; Palop et al., 2007; Roberson et al., 2007; Sanchez et al., 2012; Verret et al., 2012; Vossel et al., 2010).

One question that we initially addressed was to understand the role of hyperexcitability in AD and whether hyperexcitability occurs early in life, before cognitive impairment or plaque and tangle pathology develop. Early hyperexcitability could play a potentially more critical role than hyperexcitability later in life because it might contribute to abnormalities that develop later in life. To address this possibility, a mouse model with a slow development of impairments is useful, so that what occurs first vs. second can be discriminated. Although all mice have a rapid progression relative to humans, one model with a relatively slow progression is the Tg2576 mouse (Hsiao et al., 1996). The Tg2576 mice overexpress the Swedish mutation (APPSwe) of human amyloid precursor protein (hAPP), and amyloid-beta oligomerization occurs starting at approximately 6 months of age (Citron et al., 1992; Hsiao et al., 1996; Jacobsen et al., 2006; Kawarabayashi et al., 2001). These animals do not exhibit tangles but are nevertheless widely used to gain insight into AD. Interestingly, some types of spatial memory (e.g. novel object location) are impaired as early as 2-4 months (Duffy et al., 2015). As early as 1 month of age there is hyperexcitability, manifested by spontaneous generalized spikes in the EEG that are large in amplitude and brief (<0.5 ms) (Kam et al., 2016). Although seizures were not detected at 1-2 months of age, they did occur later in life (Kam et al., 2016). The spikes were similar to those spikes occurring between seizures, so the spikes were termed interictal (between seizures) spikes (IIS). Another group also found that IIS occur as early as 6 weeks of age in Tg2576 mice (Bezzina et al., 2015), and it has been known for many years that mice with hAPP mutations exhibit IIS although early ages may not be noted (Born et al., 2014; Palop et al., 2007; Reyes-Marin and Nunez, 2017; Sanchez et al., 2012).

IIS in Tg2576 mice appeared to be due in part to excessive activity in the cholinergic system because IIS were reduced by atropine, a muscarinic acetylcholine receptor antagonist (Kam et al., 2016). Additional approaches suggested the cholinergic projection from medial septum had increased activity and increased expression of the rate-limiting enzyme for acetylcholine synthesis, the acetylcholinesterase (Kam et al., 2016). The data suggested it was timely to reconsider the well-established idea that the medial septal cholinergic neurons play a critical role in AD (Davies and Maloney, 1976; Ferreira-Vieira et al., 2016; Mufson et al., 2008; Parent et al., 2013; Shiozaki et al., 2001; Stepanichev, 2017; Sultzer, 2018). That is, early in life in Tg2576 mice there was increased excitability manifested by IIS, and cholinergic dysfunction contributes (Kam et al., 2016). These findings are consistent with an early decrease in cholinergic markers in AD (Apelt et al., 2002; Klingner et al., 2003) and could ultimately lead to cholinergic dysfunction (Mufson et al., 2008), consistent with the cholinergic hypothesis of AD from the 1970’s (Davies and Maloney, 1976).

To define mechanisms of hyperexcitability and the role of the cholinergic neurons further, it seems useful to focus attention on the DG for several reasons. First, several neuroanatomical changes occur in the DG in hAPP mouse models (Bearer et al., 2018; Fontana et al., 2017; Jacobsen et al., 2006; Krezymon et al., 2013; Ohm, 2007; Palop et al., 2005; Palop et al., 2007; Palop et al., 2003; Roberson et al., 2011; You et al., 2017) that also occur in human AD (Ohm, 2007; Scharfman, 2012; Scharfman, 2019). Regarding hyperexcitability, the DG is relevant because it is considered to be a region that can generate seizures in temporal lobe epilepsy (Heinemann et al., 1992; Kobayashi and Buckmaster, 2003; Krook-Magnuson et al., 2015; Lothman et al., 1992; Scharfman, 2019; Sun et al., 2007). In amnestic mild cognitive impairment, neuroimaging shows that the DG and area CA3 are hyperexcitable (Bakker et al., 2012). The DG also seems to be a good choice to focus attention because of prior studies of a commonly used hAPP mouse model of AD, the J20 mouse. These mice share the APP mutation (KM670/671NL; APP_Swe_) with Tg2576 mice, as well as a second mutation (V717F; APP_Ind_) and a different promoter (platelet-derived growth factor; PDGF). This model exhibited increased expression of GC c-Fos and ΔFosB, which are upregulated in young GCs by neuronal activity (You et al., 2017).Furthermore, experimental manipulations which affected the DG led to improvement in excitability, behavior, and other measurements of dysfunction (Verret et al., 2012; You et al., 2017). Regarding the cholinergic system, the DG is also of interest because the septocholinergic neurons innervate all cell types in a robust manner (Aznavour et al., 2005; Clarke, 1985; Deller et al., 1999; Dougherty and Milner, 1999; Frotscher, 1991; Frotscher and Leranth, 1985; Frotscher and Leranth, 1986; Leranth and Frotscher, 1987; Milner and Veznedaroglu, 1993; Nyakas et al., 1987; Takacs et al., 2018; Wainer et al., 1985).

For all these reasons, we focused on the DG to obtain insights into why hyperexcitability may occur in young Tg2576 mice and the role of cholinergic input. We recorded from the principal cells of the DG (granule cells; GCs) in hippocampal slices with whole-cell recordings to understand cellular changes with high resolution. Early ages (less than 3 months) were used because of the hypothesis that at an early age, when hyperexcitability appears to occur relatively selectively, one might understand pathology that was independent of problems occurring later in life. We studied synaptic events by recording spontaneous excitatory postsynaptic potentials (sEPSPs) / currents (sEPSCs) and spontaneous inhibitory postsynaptic currents (sIPSCs). To obtain an understanding of nonsynaptic effects, intrinsic properties (e.g., resting membrane potential or RMP, input resistance or R_in_, time constant or τ, etc.) were also analyzed. Finally, we used the muscarinic antagonist atropine to determine if impairments in Tg2576 mice could be reduced by atropine. One of the reasons atropine was important to use was that the effects of atropine *in vivo* may have been due to effects on the periphery rather than the brain (Kam et al., 2016). One of the predictions of the present study was that there would be identifiable impairments in GCs in young Tg2576 mice that could help understand mechanisms of hyperexcitability observed *in vivo*.

The results suggest that there are alterations in GCs that could explain hyperexcitability as well as the alterations in the cholinergic system. The data suggest that very early (< 3 months) increases in synaptic excitation and decreased inhibition occur in GCs of Tg2576 mice. These data suggest that synaptic changes, specifically increased glutamatergic excitation and depressed GABAergic inhibition could contribute to hyperexcitability. There also were several concurrent changes in intrinsic properties that may contribute to hyperexcitability but also could do the opposite, and whether the latter is an attempt of the biological system to compensate for hyperexcitability is discussed. Finally, we obtained evidence that Tg2576 mice exhibit changes in muscarinic cholinergic regulation of GCs, some of which were novel.

Together the data provide advances in our understanding of how hyperexcitability may develop in AD. The results support a model where the earliest changes in the brain with AD are accompanied by increased synaptic excitation and decreased inhibition. Moreover, there are altered intrinsic properties and effects of muscarinic cholinergic receptors. Thus, early in life the data suggest the opposite of what is observed at late stages: instead of synaptic depression there is increased excitation. The cholinergic system is not necessarily degenerated but less able to conduct its normal function, consistent with prior views (Mufson et al., 2008). This model would be consistent with an initial overactivity early in life that has adverse effects such as inflammation, glial activation, and accumulated metabolic waste and these negative consequences contribute or even cause neuropathology and neurodegeneration.

## METHODS

### I. Animals

All experimental procedures were approved by the Institutional Animal Care and Use Committee (IACUC) at The Nathan Kline Institute and experiments were carried out in accordance with the National Institutes of Health (NIH) guidelines. Mice expressing human APP_695_ with the Swedish (Lys670Arg, Met671Leu) mutations driven by the hamster prion protein promoter (Hsiao et al, 1996) were bred from male heterozygous Tg2576 and female non-transgenic mice (C57BL6/SJL F1 hybrid, stock No. 100012, Jackson Labs), and fed a chow that is commonly used during breeding (Purina 5008, W.F. Fisher). Mice were housed with same-sex siblings and fed another common rodent food after weaning (Purina 5001, W.F. Fisher) and water, available *ad libitum*, with a 12 h light-dark cycle. Before and after any animal was used, the genotypes were confirmed using an in-house protocol for detecting the APP_695_ gene. Both male and female mice were used to determine if there were sex differences.

### II. Slice electrophysiology

#### A. Slice preparation

Mice were deeply anesthetized by isoflurane (1-2%; Aerrane, Piramal Enterprises) inhalation, followed by a subsequent intraperitoneal injection of urethane (2.5 g/kg; i.p.). Mice were perfused transcardially with a cold (4°C) sucrose-based artificial cerebrospinal fluid (sucrose ACSF) containing (in mM) 90 sucrose, 2.5 KCl, 1.25 NaH_2_PO_4_, 4.5 MgSO_4_, 25.0 NaHCO_3_, 10.0 D-glucose, 80.0 NaCl, and 0.5 CaCl_2_; pH 7.4. Then mice were decapitated and the brain was removed and dissected in the same sucrose ACSF at 4°C. All ACSF solutions were aerated with carbogen (95% O_2_, 5% CO_2_, All-Weld Products). One cerebral hemisphere was mounted on a vibratome stage, and slices were cut horizontally in cold (4°C) sucrose ACSF with a vibratome (350 μm thick; No. HMV450, Micron Instruments). Slices were immediately placed in a custom-made holding chamber containing sucrose ACSF at 30°C for 30 min and aerated with carbogen (95% O_2_, 5% CO_2_). Then, the slices were stored in the same holding chamber at room temperature for at least 60 min before recording.

#### B. Whole-cell patch clamp recordings

##### 1) Recording conditions

Slices were transferred to a recording chamber (RC-27LD, Warner) and perfused with ACSF containing NaCl instead of sucrose (NaCl ACSF) which contained (in mM: 130 NaCl, 2.5 KCl, 1.25 NaH_2_PO_4_, 1 MgSO_4_, 25.0 NaHCO_3_, 10.0 D-glucose, and 2.4 CaCl_2_ (pH 7.4). NaCl ACSF was perfused at 6 mL/min with a peristaltic pump (Masterflex C/L, Cole-Parmer) and maintained at 32°C with a temperature controller (TC-324B, Warner) and in-line heater (SH-27B, Warner).

##### 2) Recording electrodes and acquisition

For whole cell current-clamp experiments, borosilicate glass capillaries (1.5 mm outside diameter; 0.86 inner diameter, Sutter Instruments) were pulled horizontally (P-97, Sutter) so that the resistance was 4-9 MΩ. Recordings were performed with a K-gluconate based internal solution of the following composition (in mM): 130.0 K-gluconate, 4.0 KCl, 2.0 NaCl, 10.0 HEPES, 0.2 EGTA, 4.0 Mg-ATP, 0.3 Na_2_-GTP, 14.0 Tris-phosphocreatine, and 0.5% biocytin (pH of 7.25 and 302 ± 5 mOsm). Seal resistances were > 1 GΩ before breaking into whole cell configuration. All data were digitized (Digidata 1440A, Molecular Devices), amplified by a MultiClamp 700B amplifier (Molecular Devices), and low pass filtered using a single-pole RC filter at 10 kHz. Analysis used pClamp software (v11.1, Molecular Devices) and is described further below.

##### 3) Synaptic potentials and synaptic currents

For the evaluation of the synaptic activity, spontaneous events were evaluated using a continuous recording for 3-5 min.

Current-clamp recordings of spontaneous excitatory postsynaptic potentials (sEPSPs) were obtained from GCs using an intracellular solution of the following composition (in mM): K-gluconate 130, NaCl 2, HEPES 10, EGTA 0.2, Mg-ATP 4, Na-GTP 0.3, Na_2_-phosphocreatine 14, and 0.5% biocytin; adjusted to pH 7.3 and 304 mOsm. Cells were patched-clamped using the whole-cell recording configuration and sEPSPs were evaluated at the resting membrane potential of the cell.

Voltage-clamp recordings of spontaneous excitatory and inhibitory postsynaptic currents (sEPSCs and sIPSCs, respectively) were performed in a different set of neurons, using an intracellular solution of the following composition (in mM): Cesium methanesulfonate 125, NaCl 4, HEPES 10, EGTA 1, MgATP 4, Tris-GTP 0.3, diTris-phosphocreatine 10, QX-314Cl 5, and 0.2% biocytin adjusted to pH 7.3 and 290 mOsm. To evaluate sEPSCs and sIPSCs, cells were voltage-clamped at a holding potential (HP) of −70 mV and 0 mV, respectively, in order to isolate glutamatergic and GABAergic receptor-mediated currents.

Detection of spontaneous events (EPSP, EPSC and IPSC) was performed off-line using MiniAnalysis 6.0 (Synaptosoft, Inc.). sEPSPs were identified as events having a fast rise time and a peak that was >2-3 standard deviations (SD) from the root mean square (RMS) of the baseline noise (random, background electrical fluctuations) (Hong and Rebec, 2012; Serletis et al., 2011). sEPSCs and sIPSCs were included if the maximum (peak) change in current (negative or positive, respectively) relative to baseline noise had a peak amplitude of >3 SD from the RMS of the noise. This determination was chosen because pilot work showed that discrimination between synaptic events and noise was poor with 2 SD and good with >3 SD. The mean frequency and amplitude were calculated for all events over the entire recording period (each period lasted 3-5 min) and are presented as mean ± SEM.

##### 4) Intrinsic properties

For intrinsic properties, Supplementary Figure 1 shows how measurements were made. The resting membrane potential (RMP) was defined as the difference between the potential while intracellular and that recorded after withdrawing the microelectrode from the cell. Sub-threshold depolarizing and hyperpolarizing pulses (−30 pA to +30 pA; 5-10 pA steps; 1s duration) were delivered to assess input resistance (R_in_). The steady-state voltage responses were plotted against the amplitude of current injection and the slope of the linear fit between −5 pA to −30 pA was used to define R_in_ (Clampfit v. 11.1; Molecular Devices). Time constant (tau; τ) was determined from hyperpolarizing pulses (−20 pA). τ was defined as the time to reach 63% of the steady-state response. To determine characteristics of action potential (AP) discharge, an AP was used close to its threshold. Specifically, depolarizing current pulses of +40 to +100 pA (10 pA steps; 1 sec duration) were delivered to elicit an AP in approximately 50% of trials. We then used a combination of the AP analysis tool and statistics using Clampfit (pClamp software v11.1, Molecular Devices) to determine the characteristics of the APs. Threshold was defined as the membrane potential at which the AP was initiated. This was determined in Clampfit by finding the intersection of the rising phase of the AP and the slope leading up to its initiation. Then a three-point tangent slope vector was used to find the position in the initial region where the slope was at or above 10 V/s. The mean AP peak amplitude was determined from measuring RMP to the AP peak. The time to the AP peak amplitude was the time from the start of the current step to the AP peak. The time to peak from threshold involved only the period from the point of initiation of the AP to the AP peak. Half-width was defined as the time from the point of initiation to the point when the AP reached half of its peak amplitude. AP rising and decay slopes were defined by the maximum dv/dt of each AP phase, and the dv/dt ratio was defined as the ratio of rising/decay slopes. Afterhyperpolarization (AHP) amplitude was measured from the membrane potential where the AP returned to baseline to the peak of the AHP. To quantify spike frequency adaptation (Figure 7) we evaluated the time from one AP peak to the next for all the spike pairs in trains of 4 and 6-7 APs. When the train had 7 APs, only the first 6 were used.

### III. Pharmacology

All chemicals used for making the different slicing, recording and internal solutions were American chemical society (ACS) reagent grade. Atropine sulfate salt monohydrate (No. A0257), (+)-Bicuculline (No. 14340) and DL-2-amino-5-phosphonopentanoinc acid (APV; No. A5282) were obtained from Sigma-Aldrich; 6,7-dinitroquinoxaline-2,3(1H,4H)-dione disodium salt (DNQX; No. 2312) was obtained from Tocris.

Atropine was dissolved in double distilled water and a stock solution at a concentration of 10 mM was prepared. Stock solution was stored at 4°C and protected from light. For bicuculine, APV and DNQX, drugs were dissolved in double distilled water and stock solutions of every drug were prepared at the following concentrations: bicuculine 10 mM, APV 10 mM and DNQX 50 mM. Then, solutions were stored at 4°C and protected from light.

To test the effects of drugs, all recordings were performed initially at resting potential for 3-5 min in voltage- or current-clamp configurations (choice of voltage- or current-clamp are shown in the timelines for the Figures). For experiments using atropine, 10 μM was added to the ACSF and an additional period of 3-5 min was recorded after waiting for the drug to equilibrate in the recording chamber (~5 min).

For experiments using sequential addition of bicuculline (10 μM), DNQX (10 μM) and APV (50 μM) to the ACSF, there was a slightly different approach. A period of 3-5 min was recorded from the moment the drug reached the chamber. The reason for this is that the recordings showed a rapid effect of drugs which plateaued by the end of the recording period. Data were selected from the end of the recording period.

### IV. Granule cell identification

After the completion of recordings, slices were immediately placed in 4% paraformaldehyde (PFA; Sigma) in Tris buffer (TB; 0.1 M, pH 7.4) and kept in fixative at 4°C. For visualization, approximately two weeks after the experiments, slices were permeabilized with 0.7% Triton X-100 in TB with gentle but continuous shaking on a rotator at room temperature for 1 h. Then slices were incubated in a 0.1% hydrogen peroxide (H_2_O_2_) solution in TB for 30 min in order to reduce endogenous peroxidase activity. After a series of washes in 0.25% Triton X-100 in TB at room temperature, the slices were incubated in Avidin Biotin enzyme complex (Vector ABC elite Kit PK-6100 standard) for 2 h at room temperature and then pre-incubated in a solution containing 0.5 mg/mL 3,3’-diaminobenzidine (DAB; Invitrogen) and NiCl_2_ (50 mM) dissolved in TB. This solution was pre-incubated for 30 min at room temperature and followed by another solution of DAB and H_2_O_2_ (30% weight/weight) dissolved in TB for 1-2 min. Slices were washed in TB and then passed through a graded series of glycerol (25, 40, 55 %) diluted in TB for 15 min each. A subsequent incubation in 60% 2,2’-thiodiethanol (TDE; Sigma) for 30 min was performed. Then they were immediately mounted and coverslipped with 60% TDE. Photomicrographs were made on a brightfield microscope (Model BX51; Olympus of America) equipped with a charged coupled device (CCD) camera (Retiga 2000R, Teledyne QImaging) using Image-Pro Plus software (v.7; Media Cybernetics, Inc).

### V. Statistics

A total of 35 cells from 15 Tg2576 mice, and 31 cells from 14 WT mice were used for this study. No more than three GCs, from no more than two slices per animal were used. Sometimes the sample sizes are unequal because some cells were used for recording spontaneous activity, but not intrinsic properties, and vice-versa. Statistical analyses were performed using Prism 8.3 (GraphPad). For parametric data, statistical significance for comparisons between two different groups was determined using the unpaired Student’s t-test, and a Student’s paired t-test was performed to compare the same cells before and after a treatment, or for data at two time points during the same event (e.g. train of APs). For non-parametric data, Mann-Whitney *U* test was used for unpaired analysis, and the Wilcoxon test was used for paired analysis. Cumulative distributions were analyzed using the Kolmogorov-Smirnov (KS) test. For analyzing the differences in adaptation, we used a two-way ANOVA with AP pairs and genotype as main factors, followed by Sidak’s post-hoc test for the multiple comparisons of AP pairs. Interactions between factors are not reported when they are not significant. For characteristics of GC localization, Fisher’s exact test was used to compare differences statistically. All results are presented as the mean ± standard error of the mean (SEM), and statistical significance was achieved if the p value was <0.05 (denoted on all graphs by an asterisk).

## RESULTS

### I. Comparison of WT and Tg2576 synaptic activity, intrinsic properties and firing behavior

#### A. Synaptic events

##### 1) sEPSPs in WT and Tg2576 mice

GCs were recorded in current clamp to assess sEPSPs (Figure 1A). GCs from Tg2576 mice showed a significantly higher mean frequency of sEPSPs than WT mice (Tg2576: 2.28 ± 0.19 events/sec, WT: 1.59 ± 0.09; unpaired t-test, t=3.250, df=32; p=0.001; Figure 1B1a-b, 1B2a). The representative traces in Figure 1B1 show that there was primarily an increase in small events, which lowered the mean amplitude (Figure 1B2b), whereas the differences in mean amplitudes of sEPSPs were not significantly different (Tg2576: 0.39 ± 0.03 mV, WT: 0.45 ± 0.03; unpaired t-test, t=1.407, df=32; p=0.085; Figure 1B1a-b, 1B2b). The frequency distribution of sEPSPs amplitudes (Figure 1C1) showed a large number of small events in Tg2576 GCs, like the representative examples in Figure 1B1. However, the cumulative distributions of the amplitudes showed no statistical differences (Komologorov-Smirnov test, D=0.187; p=0.567; Figure 1C2), consistent with the emphasis of the cumulative distribution on amplitude rather than frequency.

**Figure 1.**
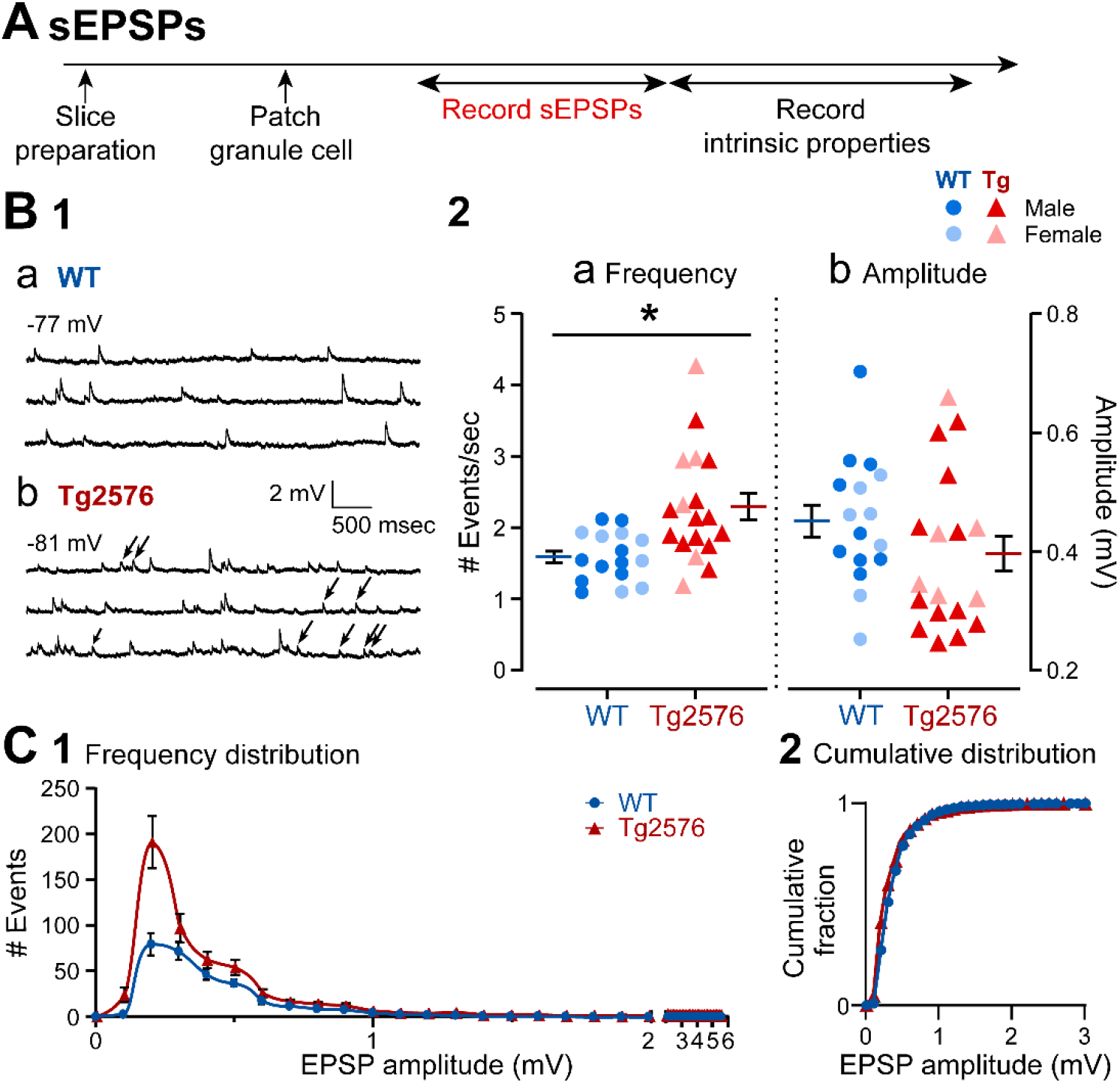
Increased frequency of sEPSPs in GCs from Tg2576 mice. (A) The timeline of the electrophysiological recordings for the determination of the sEPSPs and intrinsic properties from GCs. (B1) Representative traces show typical sEPSPs obtained in (a) WT and (b) Tg2576 mice. (B2) Quantification of sEPSP frequency and amplitude in WT and Tg2576 GCs. SEPSP frequency was significantly greater in Tg2576 mice, but not amplitude. (C1) A histogram shows the frequency distribution of sEPSP amplitudes. (C2) The cumulative distribution is shown. Differences between WT and Tg2576 GCs were not significant. Data are represented as mean ± SEM. *p<0.05. Specific p values and statistics are in the text.

##### 2) sEPSCs in WT and Tg2576 mice

Next, sEPSCs were compared in WT and Tg2576 GCs in voltage clamp (Figure 2A). The experimental timelines for voltage clamp and current clamp experiments were similar, but different GCs were sampled. Representative traces in Figure 2B1 show that sEPSCs were fast inward currents at −70 mV holding potential. Like sEPSPs, Tg2576 GCs sEPSC frequency was greater in Tg2576 mice compared to WT GCs (Tg2576 mice: 4.07 ± 0.28 events/sec, WT: 3.16 ± 0.33; unpaired t-test, t=1.998, df=18; p=0.031; Figure 2B1, 2B2a). Also, like sEPSPs, there was a lower mean amplitude of sEPSCs in Tg2576 mice. However, unlike sEPSPs, the difference was significant (Tg2576: 6.27 ± 0.54 pA; WT: 8.72 ± 1.03; unpaired t-test, t=2.432, df=18; p=0.013; Figure 2B2b). The reason why voltage clamp data were significant and current clamp data were not may be that sometimes there is a greater sensitivity in assessing synaptic events in voltage clamp compared to current clamp.

**Figure 2.**
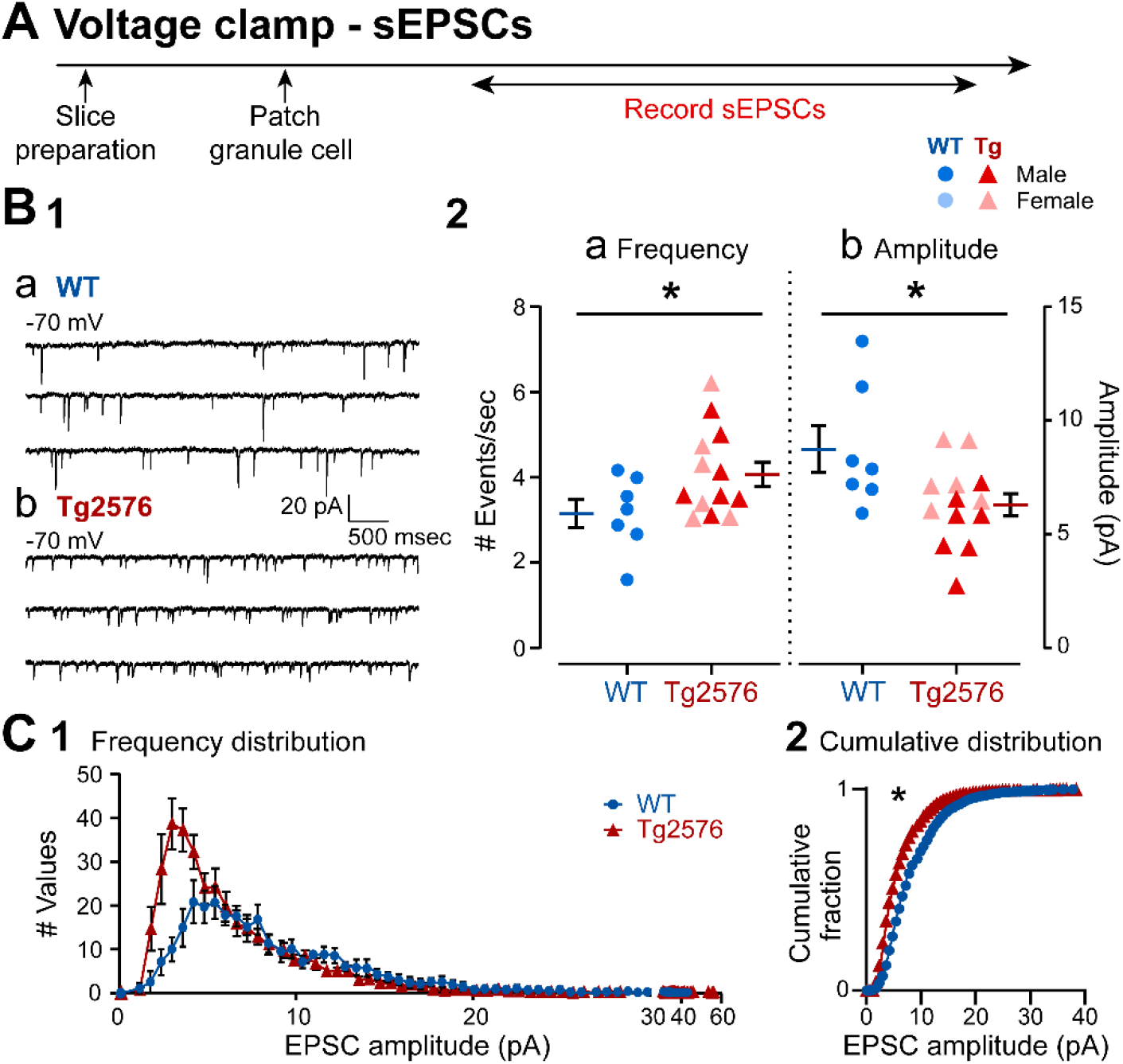
Synaptic changes in the sEPSCs in GCs from Tg2576 mice. (A) The timeline of the electrophysiological recordings for the determination of sEPSCs from GCs. (B1) Representative traces show the typical sEPSCs obtained in (a) WT and (b) Tg2576 mice from −70 mV holding potential. (B2) Quantification of sEPSC frequency and amplitude in WT and Tg2576 GCs. Tg2576 GCs had a significantly greater sEPSC frequency and a reduction in amplitude compared to WT GCs. (C1) A histogram shows the frequency distribution of sEPSCs amplitudes. (C2) The cumulative distribution is shown. The distributions of Tg2576 GCs were significantly different from WT GCs.

The frequency distribution of sEPSC amplitudes shown in Figure 2C1 illustrates a greater frequency of small sEPSCs, like data from sEPSPs in Figure 1. There was a significant difference in the cumulative distributions (Kolmogorov-Smirnov test, D=0.261, p=0.028; Figure 2C2), consistent with the data showing a significant reduction in sEPSC mean amplitudes (Figure 2B2b). Taken together, the data from current and voltage clamp suggest sEPSP/Cs increased in frequency, probably because of an increase in small sEPSP/Cs in Tg2576 mice. These small sEPSP/Cs led to a reduced mean amplitude.

##### 3) sIPSCs in WT and Tg2576 mice

The next experiments were conducted to determine if sIPSCs were different in WT and Tg2576 mice. The timeline (Figure 3A) and GCs that were sampled were the same as those used to record sEPSCs (Figure 2). The data in Figure 3B1 show that sIPSCs were recorded as fast outward currents at 0 mV holding potential. In Tg2576 mice, sIPSC frequency was significantly lower than WT mice (Tg2576: 8.19 ± 0.63 events/sec, WT: 10.38 ± 1.01; unpaired t-test, t=1.862, df=25; p=0.037, Figure 3B1, 3B2a). There were no significant differences in sIPSC mean amplitude (Tg2576: 10.57 ± 0.88 pA; WT: 10.90 ± 1.11; unpaired t-test, t=0.238, df=25; p=0.407; Figure 3B1, 3B2b). The frequency distribution of sIPSCs amplitudes in Tg2576 mice is consistent with a reduced number of events in Tg2576 mice, particularly large (>7 pA) events (Figure 3C1). There was a significant difference in the cumulative distributions (Komologorov-Smirnov test; D=0.267; p=0.002; Figure 3C2), despite the lack of change in sIPSC mean amplitudes (Figure 3B2b). The reason why the WT and Tg2576 GCs were not different in mean amplitude but cumulative distributions were different may be due to the fact that mean and cumulative distribution are not the same: cumulative distributions are more sensitive to the numbers of events of all amplitudes whereas mean amplitude pools all data.

**Figure 3.**
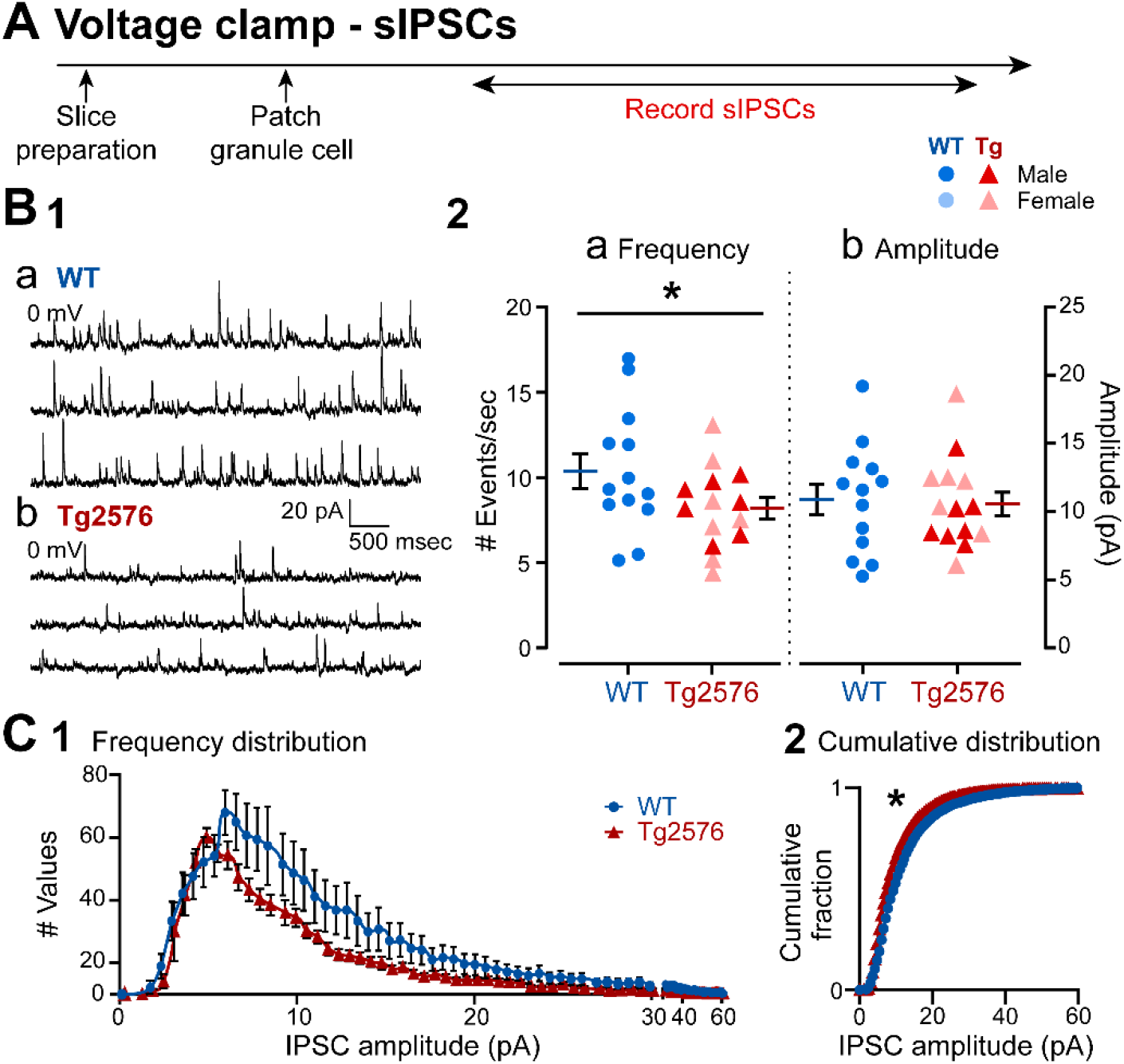
Synaptic changes in the sIPSCs in GCs from Tg2576 mice. (A) The timeline of the electrophysiological recordings for the determination of the sIPSCs from GCs. (B1) Representative traces show typical sIPSCs obtained in (a) WT and (b) Tg2576 mice at 0 mV holding potential. (B2) Quantification of sIPSC frequency and amplitude in WT and Tg2576 GCs. Tg2576 GCs had a significantly reduced sIPSC frequency, but not in amplitude compared to WT GCs. (C1) A histogram shows the frequency distribution of sIPSC amplitudes. (C2) The cumulative distributions for WT and Tg2576 GCs were significantly different.

Taken together, results in Figures 1–3 suggest increased excitatory and decreased inhibitory synaptic input to GCs of Tg2576 mice. These data suggest an altered excitatory:inhibitory (E:I) balance. The implications are important: the alterations in E:I balance in Tg2576 GCs could contribute to the increased excitability in Tg2576 mice.

#### B. Intrinsic properties in WT and Tg2576 mice

Next, we examined intrinsic properties to determine if they showed genotypic differences, and they could potentially contribute to the differences in the synaptic events of WT and Tg2576 mice (Figure 4A). The intrinsic properties that were measured were RMP, input resistance (R_in_), time constant (τ) and characteristics of APs. The characteristics of APs included amplitude, measurements of duration (time to peak, half-width), threshold, maximum rate of rise, maximum rate of decay, and dv/dt ratio (maximum rate of rise/maximum rate of decay), and the AHP amplitude (see Methods and Supplementary Figure 1).

**Figure 4.**
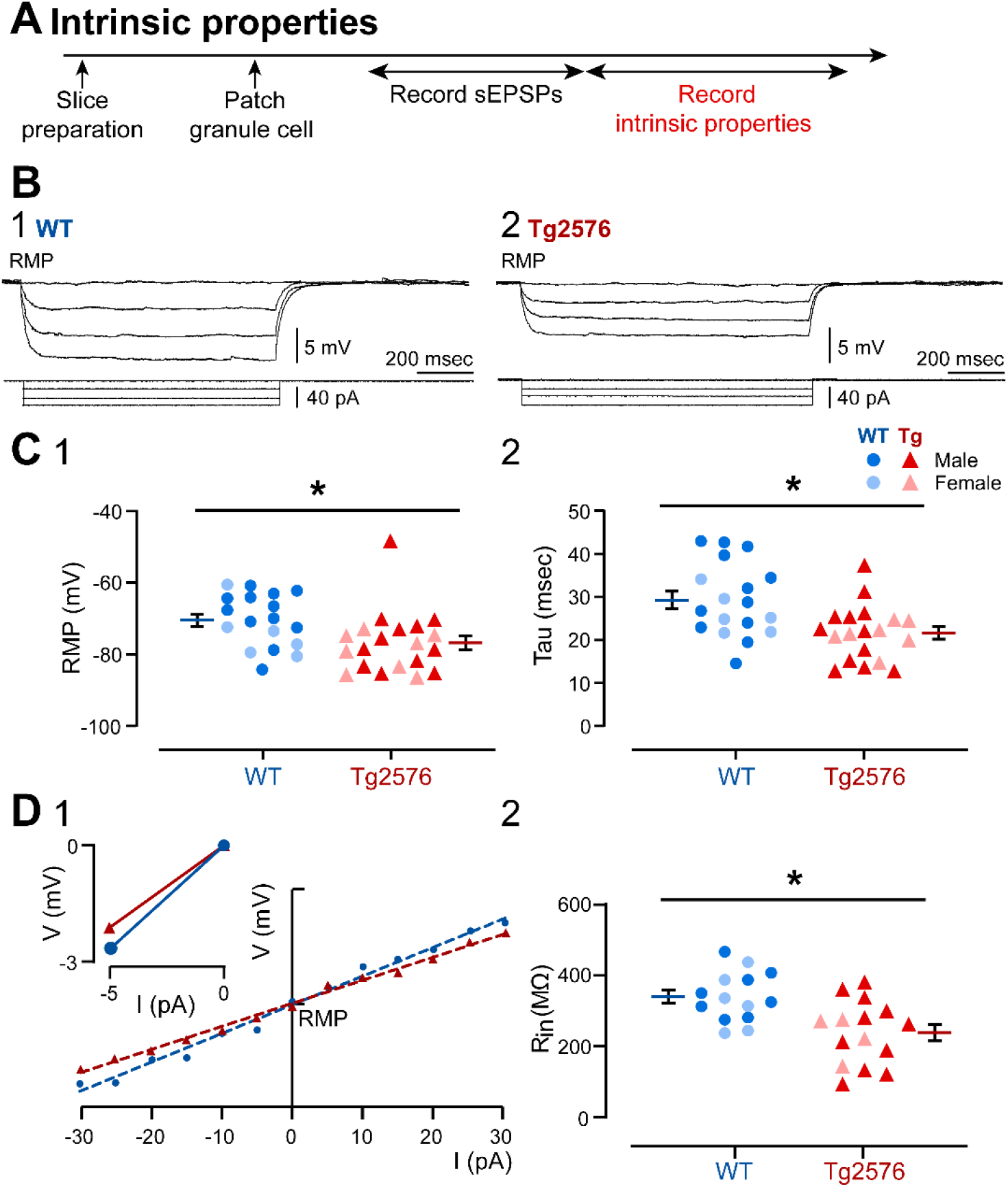
Changes in intrinsic properties of GCs from Tg2576 mice suggest a compensatory inhibitory response. (A) The timeline of the electrophysiological recordings for the determination of the intrinsic properties from GCs. (B) Membrane potential responses of representative (1) WT and (2) Tg2576 GCs to consecutive negative current steps from −5 pA to −15 pA using a 5 pA increment. (C) (1) Resting membrane potential (RMP, in mV) and (2) tau (msec) values show that Tg2576 GCs had significantly more hyperpolarized RMPs and shorter tau than WT GCs. (D1) I-V curves and linear regressions used to calculate (input resistance (R_in_). I-V curves were based on responses to positive and negative current pulses (−30 to 30 pA) in WT and Tg2576 GCs. The inset shows a representative response to a −5 pA current step from RMP, from which slope was calculated to provide a second estimation of R_in_. (D2) Average changes in R_in_ (MΩ) show a significantly lower R_in_ in Tg2576 GCs compared to WT.

The RMP of GCs from Tg2576 mice was more hyperpolarized (−77.09 ± 1.94 mV) relative to WT mice (−70.74 ± 1.75 mV; unpaired t-test, t=2.414, df=36; p=0.021, Figure 4C1). Tau was smaller in Tg2576 mice (21.46 ± 1.48 msec) than WT mice (29.17 ± 2.02 msec; unpaired t-test, t=3.108, df=35; p=0.004; Figure 4C2, 4B2). Tg2576 GCs showed a significantly smaller Rin (239.1 ± 23.07 MΩ) than WT mice (340.1 ± 18.70 MΩ; unpaired t-test, t=3.373, df=27; p=0.002; Figure 4D1, 4D2).

Representative APs and their corresponding phase plots are shown in Figure 5. Although AP amplitude and threshold were not significantly different between genotypes (Table 1), AP duration differed. The time to peak from threshold was greater in Tg2576 (1.28 ± 0.08 msec) than WT mice (1.02 ± 0.09 msec; Mann-Whitney test, U=101; p=0.032; Figure 5A and 5B). Half-width was also greater in Tg2576 (0.34 ± 0.02 msec) compared to WT mice (0.27 ± 0.02 msec; Mann-Whitney test, U=95; p=0.017; Figure 5C). Tg2576 GCs did not differ from WT mice in the maximum rate of rise, maximum rate of decay, dv/dt ratio or AHP amplitudes (Table 1).

**Figure 5.**
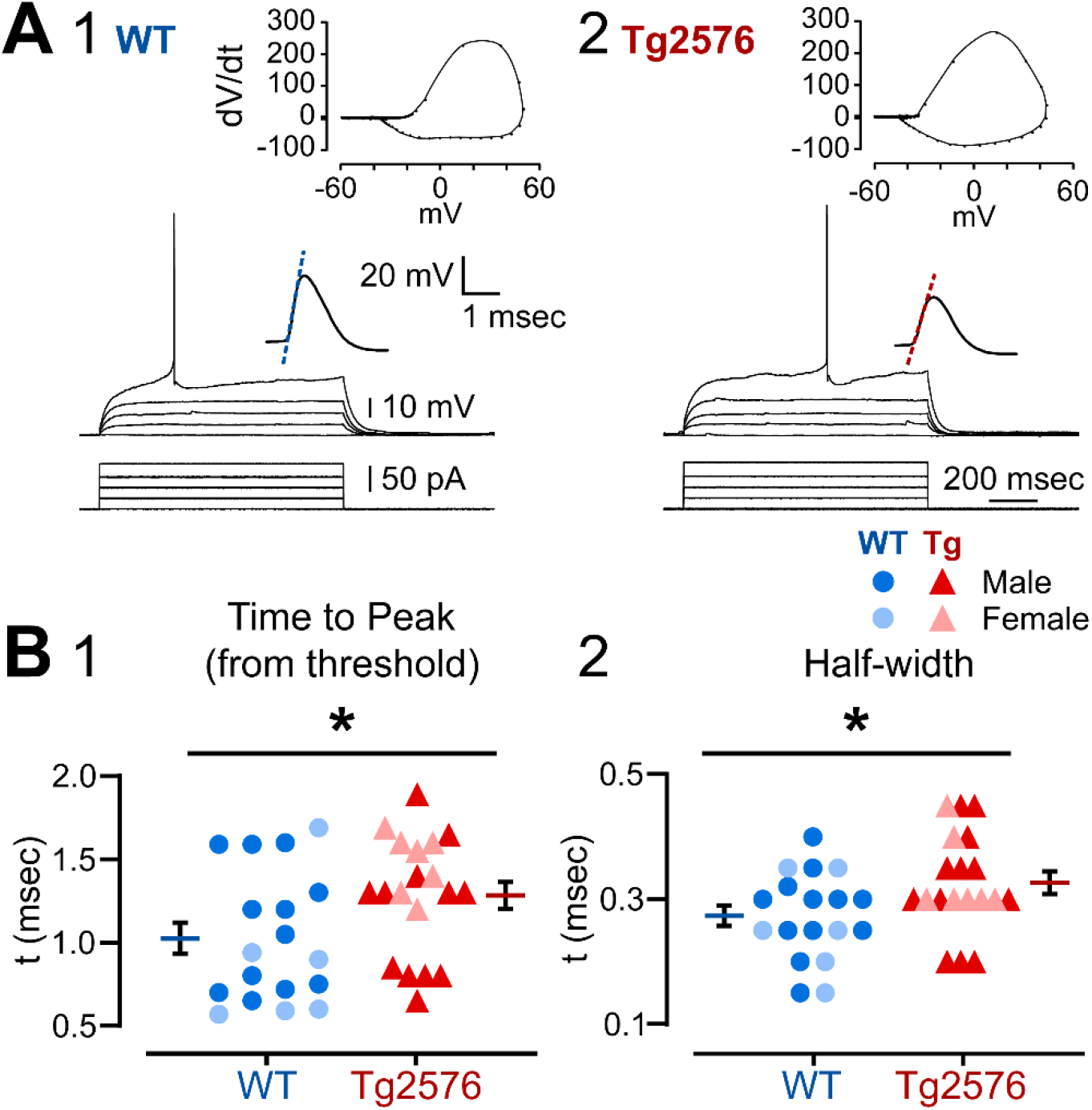
Changes in intrinsic properties of GCs from Tg2576 mice show alterations in AP properties. (A) Representative traces from (1) WT and (2) Tg2576 GCs of responses to increasingly larger current steps (+5 pA increment) from RMP until an AP was generated. The AP is shown at higher gain in the inset above the traces. The phase plots corresponding to the traces are shown at the top. (B) The mean (1) time to peak and (2) half width are shown for an AP at threshold. The APs from Tg2576 GCs had a significantly longer time to peak and half-width compared to WT GCs.

**Table 1.**
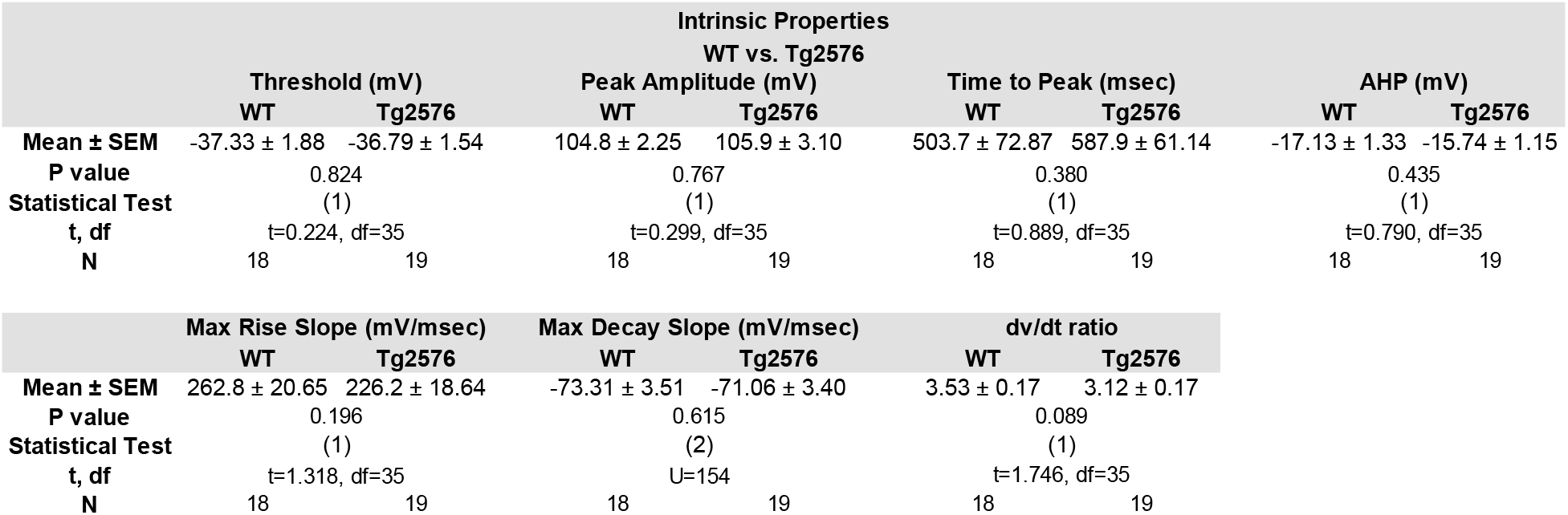
Intrinsic properties of WT and Tg2576 GCs that were not significantly different. **Table 1 legend.** Parametric data were compared using an unpaired Student’s t-test (1). Non-parametric data were compared by a Mann-Whitney test (2). Statistical significance was set at p<0.05. Data are presented as mean ± SEM.

#### C. Firing behavior in WT and Tg2576 mice

To address firing behavior, spike frequency adaptation was chosen as a measurement (Figure 6) because the firing behavior of GCs is primarily in trains with strong spike frequency adaptation, as previously reported (Scharfman, 1992; Staley et al., 1992; Williamson and Patrylo, 2007). As shown by the examples in Figure 6A, Tg2576 GCs showed weaker adaptation than WT mice. To quantify adaptation, the interspike intervals (ISI) were compared either from trains of 4 APs (Figure 6A-B) or trains that were longer (Figure 6C-D). For the comparison of 4 APs, Tg2576 GCs appeared to have less spike frequency adaptation than WT mice (Figure 6A). However, when all ISIs were compared sequentially, there was only a trend (two-way RMANOVA; F (1, 29) = 3.742; p=0.063; Figure 6B1). On the other hand, Sidak’s post-hoc test showed that the first ISI was significantly longer in Tg2576 mice (mean difference −46.98 msec; t=2.842; df=27.37; p=0.025) although other ISIs were not (all p values > 0.05; Figure 6B1).

**Figure 6.**
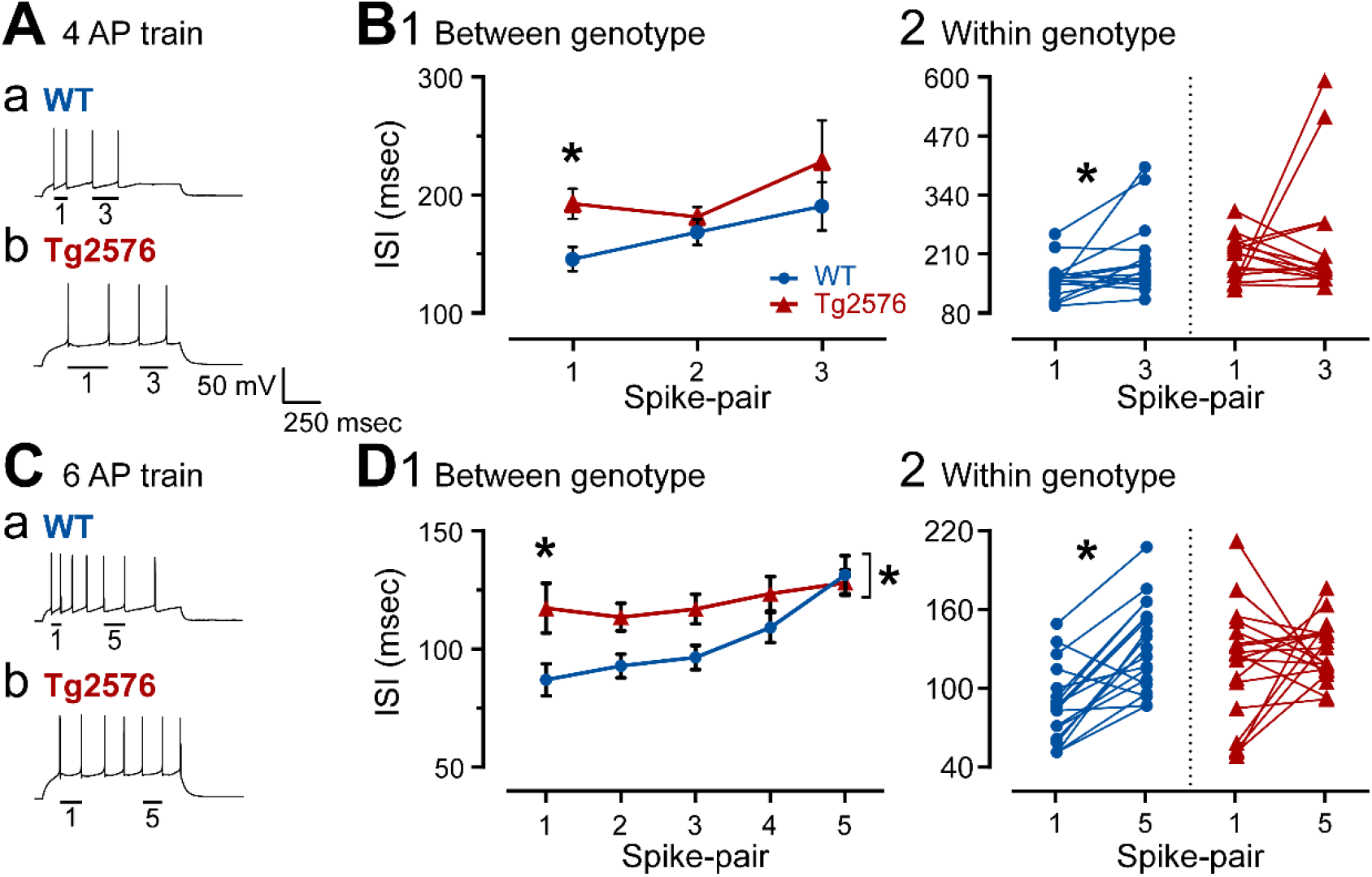
AP adaptation mechanisms are affected in GCs from Tg2576 mice. (A) Representative traces of 4 AP trains induced by positive current steps to GCs from (a) WT and (b) Tg2576 mice. The first (1) and the third (3) AP pairs are labeled by the bars below them. (B1) Spike frequency adaptation was quantified in WT and Tg2576 mice by measuring the ISI for each AP pair and plotting them sequentially. The first ISI was significantly longer ISI in Tg2576 mice. (B2) Comparisons of the first and third AP pairs showed that the first ISI was significantly shorter in WT mice, indicating adaptation occurred, but not in Tg2576 mice, suggesting Tg2576 mice have weak adaptation. (C) Representative traces of 7 AP trains induced by positive current steps to GCs from (a) WT and (b) Tg2576 mice. The first (1) and 5th (5) AP pairs are labeled by the bars below the traces. (D1) Spike frequency adaptation was quantified in WT and Tg2576 mice by measuring the ISI for each AP pair and plotting them sequentially. The first ISI was significantly longer in Tg2576 mice. The sequence of ISIs were also significantly different, with Tg2576 mice exhibiting longer ISIs. (D2) Comparisons of the first and fifth AP pairs showed that the first ISI was significantly shorter in WT mice, indicating adaptation occurred, but not Tg2576 mice, suggesting Tg2576 mice have weak adaptation.

When the first and third AP pairs were examined specifically, the WT mice showed adaptation (first AP pair: 141.3 ± 10.39 msec; third AP pair: 186.1 ± 20.56; Wilcoxon’s test, W=92, p=0.016; Figure 6B2). In contrast, Tg2576 mice did not exhibit significant differences between the first and third pair of APs (first AP pair: 188.3 ± 12.85 msec; third AP pair: 223.7 ± 35.09; Wilcoxon test, W=-12, p=0.762, Figure 6B2).

These data suggested weak adaptation using a 4 AP train, especially in Tg2576 mice. Adaptation using longer trains (6 APs) was studied to confirm these results (Figure 6C). The sequence of APs was significantly different in WT and Tg2576 mice (two-way RMANOVA; F(1, 170)= 14.24; p=0.0002; Figure 6D1) and Sidak’s post-hoc test showed a significant difference for the first AP pair (mean difference −30.20 msec; t=3.091; df=170; p=0.012) but not the others (all p values > 0.05; Figure 6D1), similar to the 4 AP train. In Figure 6D2, comparisons were made of WT and Tg2576 GCs for the first and fifth AP pairs only. The data showed that there was adaptation in the WT GCs (first AP pair: 87.08 ± 6.91 msec; fifth AP pair: 131.5 ± 8.08; paired t-test, t=5.458, df=17; p<0.0001; Figure 6D2) but not the Tg2576 GCs (first AP pair: 117.3 ± 10.56 msec; fifth AP pair: 128.1 ± 5.40; paired t-test, t=0.814, df=17; p=0.427; Figure 6D2). These data confirm the results from 4 AP trains and suggest a potential reason that there is increased excitability in Tg2576 mice compared to WT mice: Tg2576 GCs showed weak spike frequency adaptation.

The longer AP time to peak and half-width in Tg2576 mice could potentially contribute to the greater ISI for the first AP pair of a train in Tg2576 mice. These changes could delay firing of the first AP or the second, which appeared to occur in some Tg2576 GCs (Figures 5A2, 6A and 6C). The differences in intrinsic properties could also change the ISI during AP trains in Tg2576 GCs by increasing the duration of an AP and possibly as a result, the refractory period, delaying the next AP.

The relatively weak spike frequency adaptation in Tg2576 GCs is interesting because the muscarinic cholinergic “M” current is a major contributing factor to spike frequency adaptation in prior studies of pyramidal cells (Rogawski, 2000; Storm, 1990). To understand cholinergic regulation of GCs in WT and Tg2576 mice further, we examined the role of the muscarinic cholinergic receptor on sPSCs, intrinsic properties, and firing behavior.

### II. Muscarinic cholinergic modulation of synaptic activity, intrinsic properties and firing behavior in WT vs. Tg2576 mice

#### A. Synaptic activity

##### 1) sEPSPs in WT and Tg2576 mice

To address how muscarinic cholinergic receptors might contribute to the differences between sEPSPs in WT and Tg2576 mice, the effects of the muscarinic antagonist atropine was tested by adding it to the ACSF (10 μM; Figure 7A).

**Figure 7.**
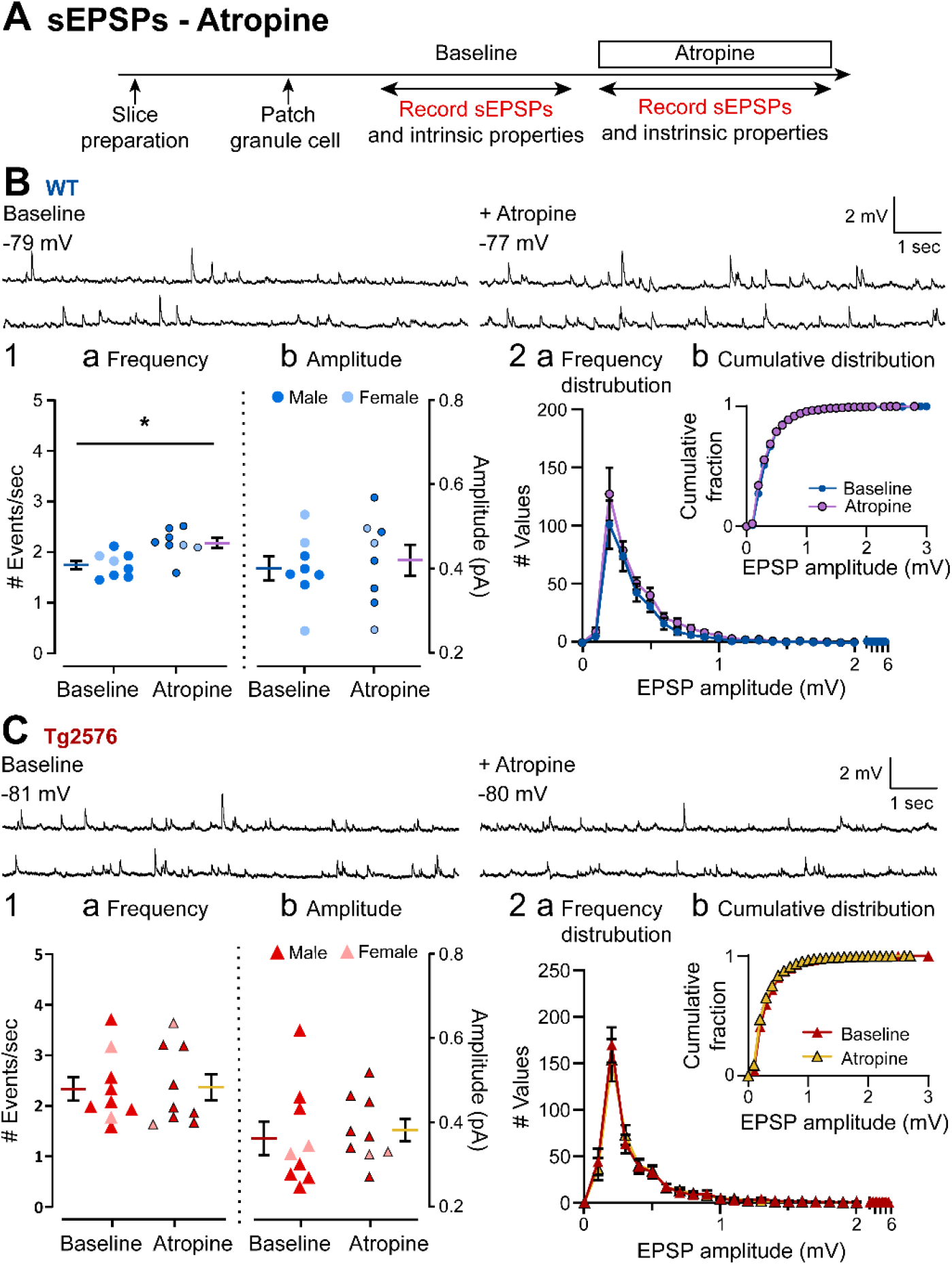
Atropine produces a small increase in sEPSP frequency in WT GCs but not in Tg2576 GCs. (A) The timeline of the electrophysiological recordings for the evaluation of the effects of atropine on sEPSPs. Atropine (10 μM) was added to the ACSF for the time period indicated by the box. (B) Representative traces for sEPSPs for WT GCs obtained in baseline conditions (left) and in presence of atropine (right). (B1) Quantification of the (a) frequency and (b) amplitude of sEPSPs for WT GCs are presented. The only effects of atropine that were significant was that atropine increased the mean sEPSP frequency. The magnitude of this effect was small. (B2) Histograms showing the (a) frequency distribution of sEPSPs amplitudes for WT GCs. Insets (b) show that there were no significant effects of atropine on cumulative distributions. (C) Representative traces for sEPSPs for Tg2576 GCs obtained in baseline conditions (left) and in presence of atropine (right). (C1) Quantification of the (a) frequency and (b) amplitude of sEPSPs for Tg2576 GCs are presented. There was no effect in Tg2576 mice. (C2) Histograms showing the (a) frequency distribution of sEPSPs amplitudes for Tg2576 GCs. Insets (b) show that there were no significant effects of atropine on cumulative distributions.

Current clamp experiments showed that WT sEPSPs increased in response to atropine, but this was not true for Tg2576 sEPSPs (Figure 7). Thus, the mean frequency of sEPSPs in WT mice was 1.75 ± 0.09 events/sec before atropine and after atropine it was 2.18 ± 0.10 (paired t-test, t=3.309, df=7, p=0.013; Figure 7B1a). In Tg2576 mice, the frequencies before and after atropine were 2.15 ± 0.23 and 2.19 ± 0.26 events/sec, respectively, which was not significantly different (paired t-test, t=0.247, df=8, p=0.811; Figure 7C1a). One reason why the Tg2576 mice did not show an effect of atropine might be that the baseline frequency of Tg2576 mice was already high (2.15 ± 0.23 events/sec) relative to WT mice (1.75 ± 0.09 events/sec) before atropine. Thus, the baseline sEPSP frequency of Tg2576 GCs (2.15 ± 0.23 events/sec) was similar to the post-atropine sEPSP frequency of WT GCs (2.18 ± 0.10 events/sec; unpaired t-test, t=0.127, df=10.89, p=0.901; Figure 7B1a vs. 7C1a).

In contrast to frequency, there was no effect of atropine on sEPSP mean amplitude in WT mice (baseline: 0.40 ± 0.03 mV; atropine: 0.42 ± 0.04; paired t-test, t=0.604, df=7; Figure 7B1b) or Tg2576 mice (baseline: 0.35 ± 0.04 mV; atropine: 0.37 ± 0.03; paired t-test, t=0.985, df=7, p=0.358; Figure 7C1b).

Consistent with a lack of significant effects of atropine on sEPSP amplitude, WT and Tg2576 mice did not show any differences in their frequency distributions (Figure 7B2, 7C2), or cumulative distributions (Kolmogorov-Smirnov tests in WT: D=0.166, p=0.826; Figure 7B2b; Tg2576: D=0.095, p=0.999; Figure 7C2b).

##### 2) sEPSCs in WT and Tg2576 mice

Next, atropine was tested using voltage clamp and sEPSCs were analyzed (Figure 8A). Atropine did not have a significant effect on sEPSC frequency in WT mice (baseline: 3.16 ± 0.33 events/sec; atropine: 3.44 ±0.33; paired t-test, t=1.221, df=6, p=0.268; Figure 8B1a) or sEPSC amplitude (baseline 8.72 ± 1.03 pA; atropine: 7.84 ± 0.88; paired t-test, t=1.656, df=6, p=0.149; Figure 8B1b). Consistent with the lack of changes in frequency and amplitude, the frequency distributions showed no differences between recordings before vs. after atropine for WT mice, and there were no significant differences in the cumulative distributions (Kolmogorov-Smirnov test, D=0.129; p=0.734; Figure 8B2b).

**Figure 8.**
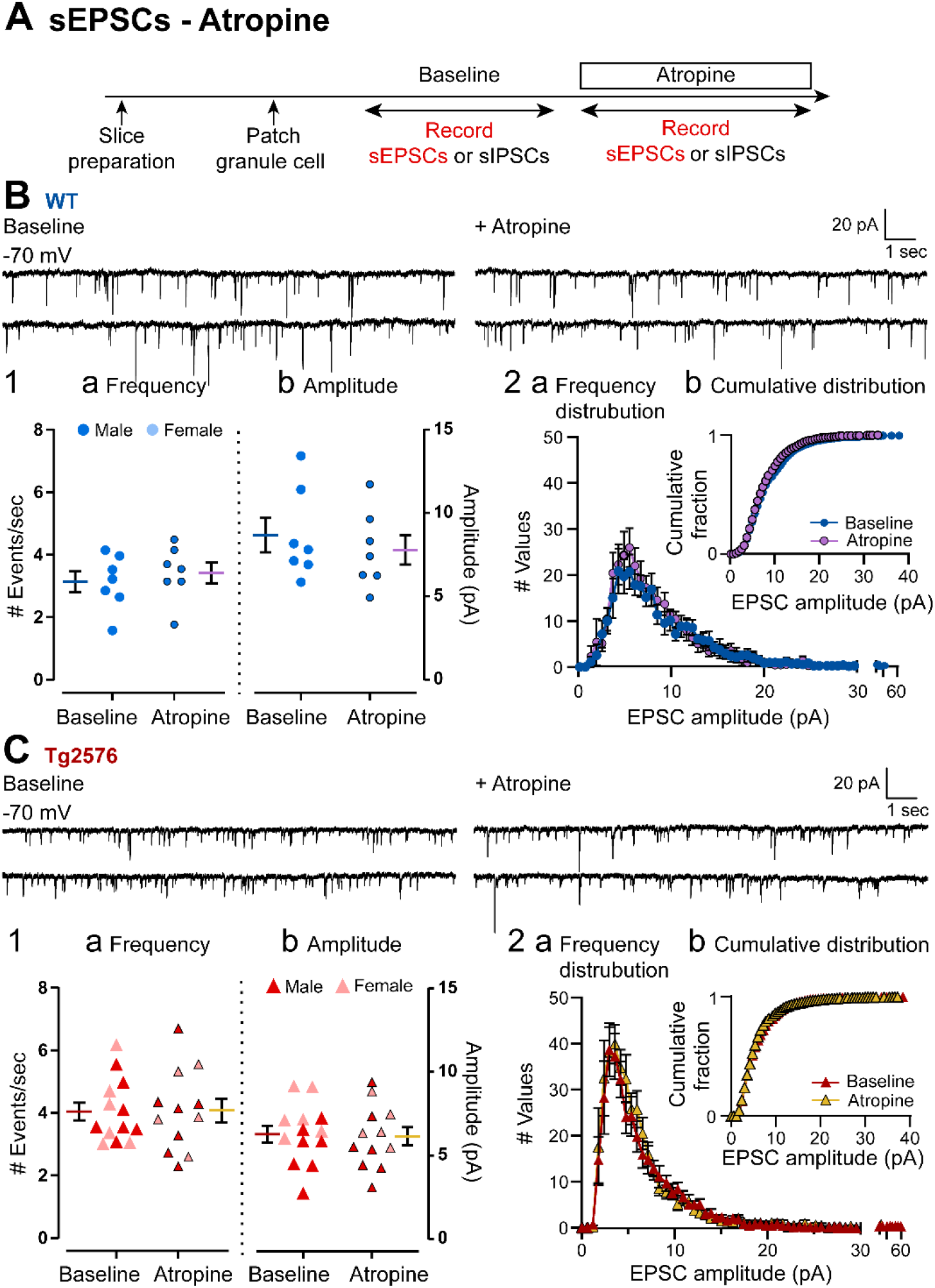
Atropine did not affect sEPSCs in GCs of WT and Tg2576 mice. (A) The timeline of the electrophysiological recordings to determine effects of atropine on sEPSCs of WT and Tg2576 GCs. (B) Representative traces for sEPSCs for WT GCs obtained in baseline conditions (left) and in presence of atropine (right). (B1) Quantification of the (a) frequency and (b) amplitude of sEPSCs for WT GCs showed no significant effects of atropine. (B2) Histograms showing the (a) frequency distribution of sEPSCs amplitudes for WT GCs. Insets (b) show that there were no significant effects of atropine on cumulative distributions. (C) Representative traces for sEPSCs for Tg2576 GCs obtained in baseline conditions (left) and in presence of atropine (right). (C1) Quantification of the (a) frequency and (b) amplitude of sEPSCs for Tg2576 GCs showed no significant effects of atropine. (C2) Histograms showing the (a) frequency distribution of sEPSCs amplitudes for Tg2576 GCs. Insets (b) show that there were no significant effects of atropine on cumulative distributions.

Next, Tg2576 sEPSCs were evaluated (Figure 8C). Like WT mice, atropine did not have a significant effect on sEPSC frequency (baseline: 4.05 ± 0.31 events/sec; atropine: 4.10 ± 0.28; paired t-test, t=0.169, df=11; p=0.870; Figure 8C1a) or sEPSC amplitude (baseline: 6.26 ± 0.54 pA; atropine: 6.15 ± 0.54; paired t-test, t=0.311, df=11, p=0.762; Figure 8C1b). Similar to WT mice, there was not a significant effect of atropine on cumulative distributions of Tg2576 mice (Kolmogorov-Smirnov test, D=0.091; p=0.963; Figure 8C2b).

##### 3) sIPSCs in WT and Tg2576 mice

Next, the effect of atropine on sIPSCs was addressed using the same experimental timeline to the one used for sEPSCs (Figure 9A). There was a significant reduction in sIPSC frequency in WT GCs (baseline: 10.24 ± 1.09 events/sec; atropine: 7.96 ± 0.86; paired t-test, t=2.456, df=11, p=0.032; Figure 9B1a) and Tg2576 GCs (baseline: 8.19 ± 0.63 events/sec; atropine: 6.42 ± 0.53; paired t-test, t=4.477, df=13, p=0.0006; Figure 9C1a). Atropine also reduced sIPSC mean amplitude in WT GCs (baseline: 10.93 ± 1.21 pA; atropine: 8.74 ± 0.76 pA; paired t-test, t=3.790, df=11, p=0.003; Figure 9B1b) and Tg2576 GCs (baseline: 10.57 ± 0.88 pA; atropine: 9.18 ± 0.74; Wilcoxon test, W=−77.0, p=0.013; Figure 9C1b).

**Figure 9.**
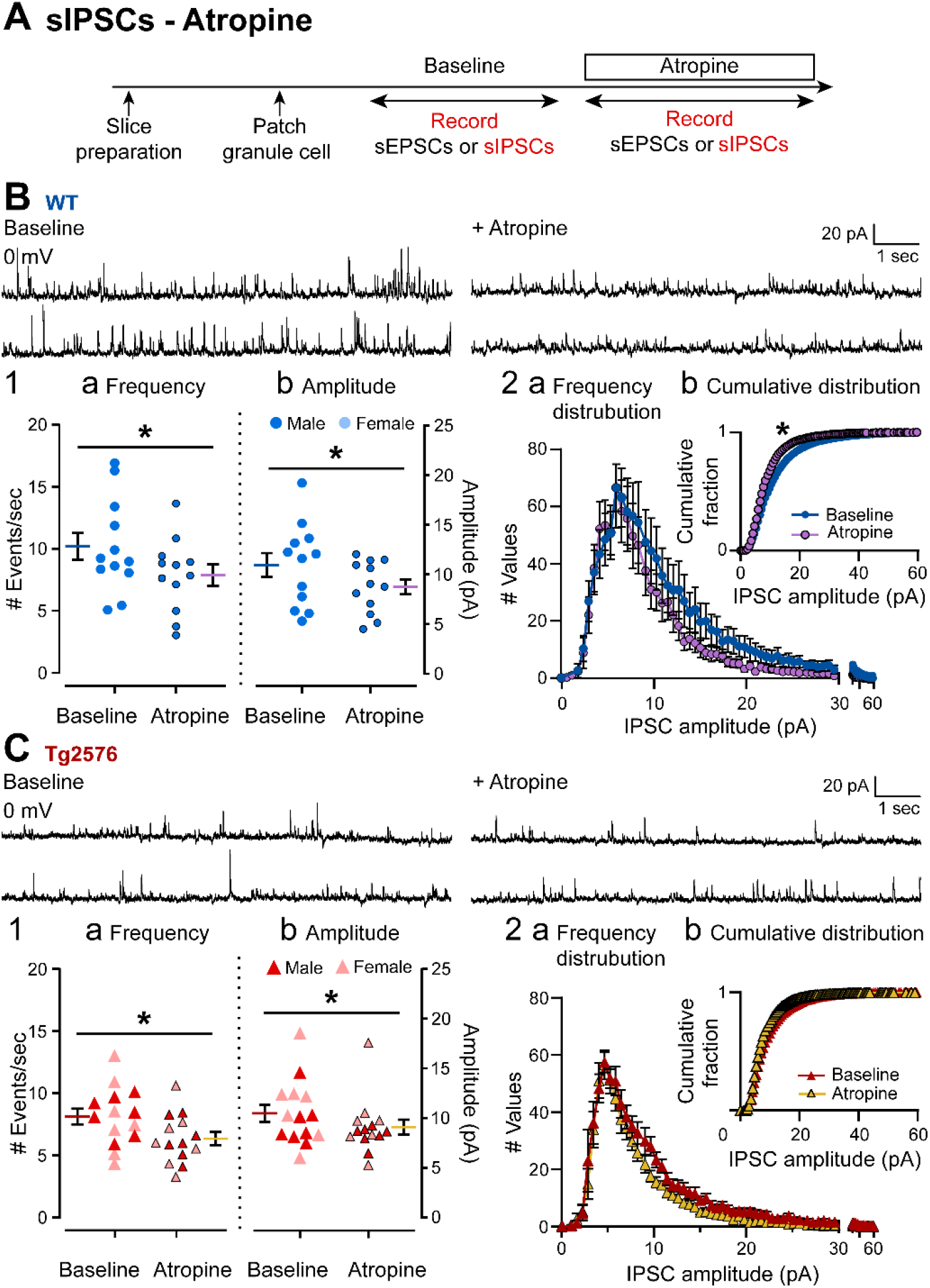
Atropine reduced sIPSC frequency and amplitude in both WT and Tg2576 mice. (A) The timeline of electrophysiological recordings used to determine effects of atropine on sIPSCs of GCs from WT and Tg2576 mice. (B) Representative traces of sIPSCs for WT GCs obtained in baseline conditions (left) and in presence of atropine (right). (B1) Quantification of the (a) frequency and (b) amplitude of sIPSCs for WT GCs show that atropine reduced frequency and amplitude in WT mice. (B2) Histograms showing the (a) frequency distribution of sIPSPs amplitudes for WT GCs. Insets (b) show the cumulative distributions where there is a significant effect of atropine in WT mice. (C) Representative traces of sIPSCs for Tg2576 GCs obtained in baseline conditions (left) and in presence of atropine (right). (C1) Quantification of the (a) frequency and (b) amplitude of sIPSCs for Tg2576 GCs show that atropine reduced frequency and amplitude in Tg2576 mice. (C2) Histograms showing the (a) frequency distribution of sIPSPs amplitudes for Tg2576 GCs. Insets (b) show the cumulative distributions. There was not a significant effect of atropine in Tg2576 mice.

The frequency distributions for WT and Tg2576 mice showed different effects of atropine (Figure 9B2, 9C2). The cumulative distributions of sIPSCs showed a decline in sIPSCs after atropine that was significant for WT mice (Kolmogorov-Smirnov test, D=0.250, p=0.006; Figure 9B2b), but not for Tg2576 mice (Kolmogorov-Smirnov test, D=0.104, p=0.740 Figure 9C2b). These data suggest that muscarinic cholinergic receptors decrease sIPSCs in WT and Tg2576 GCs.

##### 4) A prominent type of excitatory synaptic current in Tg2576 GCs and effects of atropine

In the course of the voltage clamp experiments, recordings at 0 mV revealed additional notable findings (Figure 10A, Supplemental Figure 2C). There were small inward glutamatergic synaptic currents that we defined as novel (novel spontaneous currents, nsCs) because they were especially prominent during the baseline in Tg2576 GCs (Figure 10A-D) relative to WT GCs. Atropine appeared to induce nsCs in WT mice but had less of an effect in Tg2576 mice (Figure 10A-D), perhaps because there already were many nsCs present.

**Figure 10.**
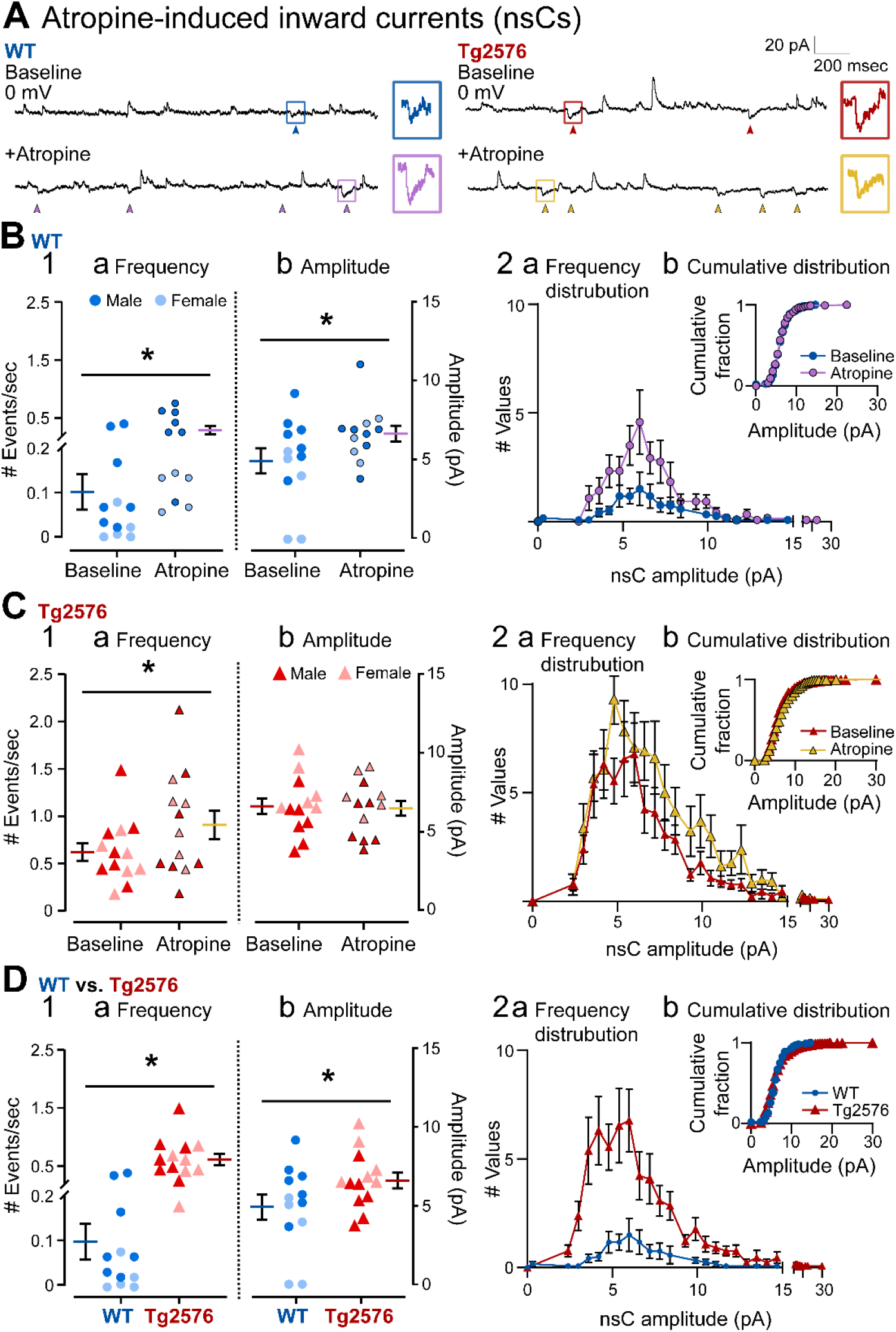
A novel spontaneous inward current (nsC) and the effects of atropine. (A) Representative traces show spontaneous inward currents (pointed with arrows) obtained in WT (left) and Tg2576 mice (right) in baseline conditions and in the presence of atropine (10 μM). Holding potential was 0 mV. The small squares mark examples of the spontaneous inward currents, which we refer to as nsCs. The examples of nsCs are expanded on the right. (B1) Mean frequency (a) and amplitude (b) of nsCs are shown for WT mice during the baseline and after atropine was added. Atropine significantly increased the frequency and amplitude of nsCs. (B2) WT nsC frequency distributions (a) and cumulative distributions (b) are shown. Atropine did not cause significant changes to the cumulative distributions relative to baseline conditions. (C1) Mean frequency (a) and amplitude (b) of nsCs are shown for Tg2576 mice. Atropine significantly increased the frequency of nsCs but not the amplitude. (C2) Tg2576 nsC frequency distributions (a) and cumulative distributions (b) are shown. Atropine did not cause significant changes to the cumulative distributions, like WT mice. (D1) Comparisons of WT and Tg2576 GCs for nsC frequency (a) and amplitude (b) during baseline conditions. Tg2576 GCs had significantly more frequent and larger nsCs than WT mice. (D2) NsC frequency distributions (a) and cumulative distributions (b) for WT and Tg2576 GCs during baseline conditions are shown. The cumulative distribution of nsCs did not show a significant difference.

We confirmed that nsCs were glutamatergic, like other inward currents at 0 mV (Supplemental Figure 2) and at the same time, confirmed that the sEPSCs described in the previous sections were glutamatergic, and sIPSCs were GABAergic (Supplemental Figure 2B and 2C). In these experiments, after atropine was added, the GABA_A_ receptor antagonist bicuculline was added also (Supplemental Figure 2A), and sIPSCs were blocked (Supplementary Figure 2B3 and 2C2). Afterwards, CNQX and APV were added, and inward currents were blocked (Supplemental Figure 2B4 and 2C3).

Because of their small size, we discriminated nsCs from the baseline noise by their kinetics (fast rate of rise relative to decay, like synaptic currents) and amplitude >1 standard deviation (SD) greater than the baseline noise (Supplemental Figure 2C1, arrows). Baseline noise was defined as the average peak-to-peak amplitude for a randomly-selected period of the baseline lasting 1 sec.

The presence of nsCs was rare in WT GCs under baseline conditions, as mentioned above. This may be why they have not been previously reported. Also mentioned above, Tg2576 GCs showed more frequent nsCs during the baseline than WT GCs. This and other direct WT vs. Tg2576 comparisons are explained further below (Figure 10D).

Effects of atropine (10 μM) on nsCs showed some similarities and some differences between WT and Tg2576 mice (Figure 10A-C). For WT nsCs, atropine increased frequency (baseline: 0.10 ± 0.04 events/sec; atropine: 0.29 ± 0.07; Wilcoxon test, W=78; p<0.001, Figure 10B1a) as well as amplitude (baseline: 4.91 ± 0.79 pA; atropine: 6.32 ± 0.51; paired t-test, t=2.302, df=11; p=0.021; Figure 10B1b).

Atropine also increased the frequency of nsCs in Tg2576 GCs (baseline: 0.63 ± 0.09 events/sec; atropine: 0.92 ± 0.15; paired t-test, t=2.758, df=12; p=0.017, Figure 10C1) However, nsC amplitude did not change significantly in response to atropine (baseline: 6.59 ± 0.50 pA; atropine: 6.46 ± 0.48; paired t-test, t=0.523, df=12; p=0.610; Figure 10C1b), probably because Tg2576 nsCs were already large in amplitude before atropine. Thus, WT nsC mean amplitude after atropine was approximately the same as the Tg2576 nsC mean amplitude before atropine.

When comparing the frequency distributions, the data reflected the larger changes after atropine in WT mice (Figure 10B2a), compared with Tg2576 mice (Figure 10C2a). Cumulative distributions did not show any significant difference before and after atropine in WT (Kolmogorov-Smirnov test, D=0.349; p=0.188; Figure 10B2b) and Tg2576 mice (Kolmogorov-Smirnov test, D=0.241; p=0.351; Figure 10C2b).

When WT and Tg2576 mice were compared directly (Figure 10D) several interesting findings were revealed. Under baseline conditions, nsCs were approximately 6-fold greater in frequency in Tg2576 mice compared to WT mice (Tg2576: 0.63 ± 0.09, events/sec; WT: 0.10 ± 0.04; Mann-Whitney test, *U*=4, p<0.001, Figure 10A and 10D1a). during the baseline, nsCs were also slightly greater in amplitude in Tg2576 mice relative to WT mice (Tg2576: 6.59 ± 0.50 pA, and WT: 4.91 ± 0.79; unpaired t-test, t=1.833, df=23; p=0.040; Figure 10A,10D1b). The frequency distributions reflecting the differences between WT and Tg2576 nsCs are shown in Figure 10D2a. The cumulative distributions were not significantly different (Kolmogorov-Smirnov test, D=0.344, p=0.131; Figure 10D2b).

Taken together, nsCs reflect a glutamatergic synaptic current that was more prevalent in Tg2576 GCs relative to WT GCs. While atropine induced more nsCs in both WT and Tg2576 mice, there were greater effects of atropine in WT mice. These data may explain the more frequent small sEPSPs in Tg2576 compared to WT mice (Figure 1B2a). The data may also reflect an alteration in the actions of muscarinic receptors in young Tg2576 mice so that effects of atropine were diminished compared to effects of atropine in WT mice.

#### B. Intrinsic properties

To determine if atropine altered intrinsic properties, GCs used to record sEPSPs were recorded before and after atropine (Figure 7A). In WT and Tg2576 mice, most intrinsic properties were not significantly different when pre-and post-atropine values were compared (Table 2). However, R_in_ was higher in Tg2576 GCs after atropine (baseline: 262.7 ± 35.64 MΩ; atropine: 451.9 ± 63.68; paired t-test, t=2.767, df=5; p=0.040; Table 2). In WT mice, atropine also increased mean R_in_ but the difference from the baseline mean was not significant (baseline: 295.3 ± 32.21 MΩ; atropine: 372.8 ± 67.72; paired t-test, t=0.891, df=3; p=0.439; Table 2). Another difference between WT and Tg2576 mice was threshold. The mean threshold was more depolarized after atropine in Tg2576 GCs, although the difference from the baseline mean was only 2 mV (baseline: −33.07 ± 2.23 mV; atropine: −31.37 ± 2.49; paired t-test, t=2.819, df=8; p=0.023; Table 2). Threshold was not significantly changed by atropine in WT mice (baseline: −39.76 ± 2.89 mV; atropine: −39.23 ± 2.95, paired t-test, t=0.951, df=7, p=0.373; Table 2). These data shed light on potential reasons for the increases in nsCs in Tg2576 mice after atropine because the increase in R_in_ in Tg2576 mice could explain an increased frequency of small synaptic events.

**Table 2.**
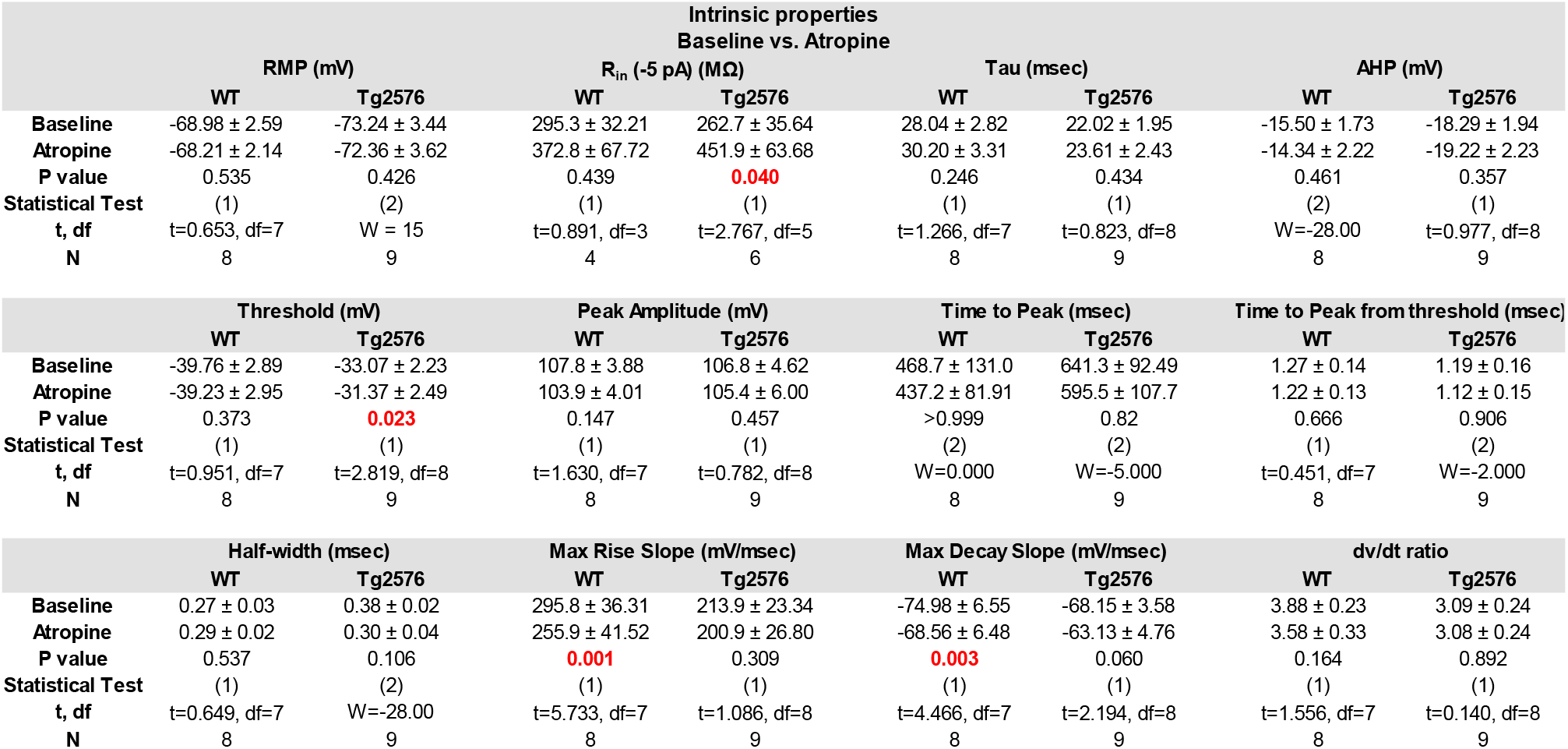
Effects of atropine on GC intrinsic properties. **Table 2 legend.** Intrinsic properties of WT and Tg2576 GCs during baseline and after atropine was added. Parametric data were compared using a paired Student’s t-test (1). Non-parametric data were compared by a Wilcoxon matched pairs test (2). Red font indicates significant differences (p<0.05).

In WT mice, the maximum rate of rise and rate of decay were decreased by atropine (maximum rate of rise, baseline: 295.8 ± 36.31 mV/msec; atropine: 255.9 ± 41.52; paired t-test, t=5.733, df=7, p=0.001; Maximum rate of decay, baseline: −74.98 ± 6.55 mV/msec; atropine: −68.56 ± 6.48, paired t-test, t=4.466, df=7, p=0.003; Table 2). However, atropine did not have a significant effect in Tg2576 mice, possibly because the rate of rise and decay were already low before atropine.

#### C. Firing behavior

Next, spike frequency adaptation was examined. When trains of 4 APs were analyzed, there were no differences between WT and Tg2576 mice in the sequence of APs compared by two-way RMANOVA (WT: F(1,21)=0.1724, p=0.682; Tg2576: F(1,20)=0.3530, p=0.559; Figure 11A-B1). When the first and third AP pairs were compared, WT mice showed adaptation during the baseline, but there was no significant effect of atropine (paired t-test, t=1.423, df=6, p=0.102; Figure 11B2). Tg2576 mice did not show adaptation and there was no significant effect of atropine (paired t-tests, t=0.671, df=6, p=0.264; Figure 11B2).

**Figure 11.**
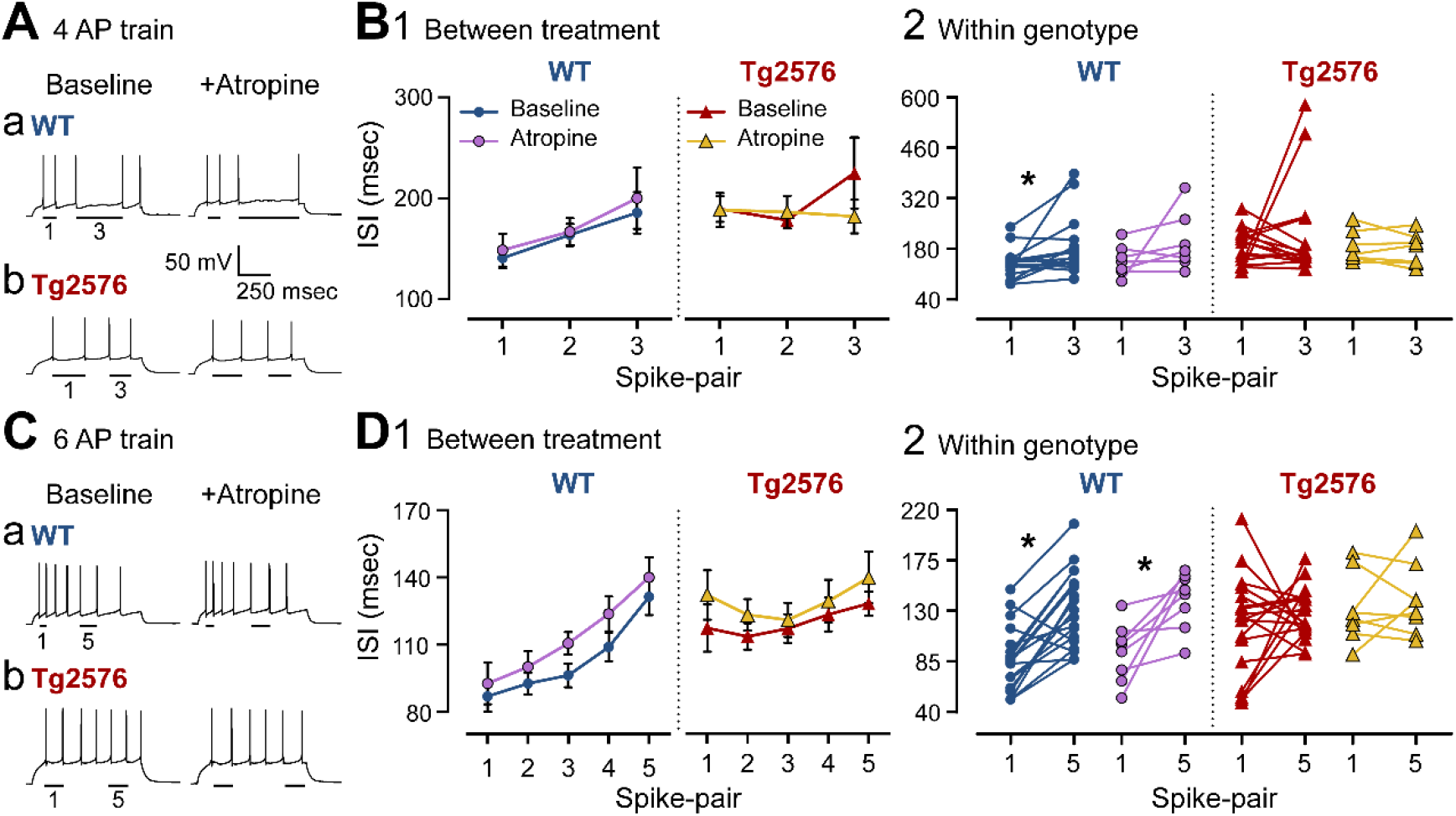
Atropine has little effect on interspike intervals of WT and Tg2576 GCs. (A) Representative traces of 4 AP trains induced by depolarizing current pulses during the baseline (left) or the presence of atropine (right). (a) WT and (b) Tg2576 GCs. (B1) Mean ISIs are plotted to show all AP pairs of the trains in WT (left) and Tg2576 GCs (right). There was spike frequency adaptation during the baseline in WT but not Tg2576 mice. Atropine did not have a significant effect. (B2) ISIs for the first and third AP pairs are plotted for WT (left) and Tg2576 mice (right) during the baseline or the presence of atropine. WT mice showed a significant increase in ISI between AP pairs during the baseline but Tg2576 mice did not. In the presence of atropine there was no longer a significant increase in the ISI of the third pair, suggesting that atropine reduced spike frequency adaptation in WT mice. In Tg2576 mice there was no significant difference in ISIs during the baseline and the same was true in the presence of atropine. (C) Representative traces of 7 AP trains induced by depolarizing current pulses during the baseline (left) or the presence of atropine (right). (a) WT and (b) Tg2576 GCs. (D1) Mean ISIs are plotted to show all AP pairs of the trains in WT (left) and Tg2576 GCs (right). There was spike frequency adaptation during the baseline in WT but not Tg2576 mice. Atropine did not have a significant effect. (D2) ISIs for the first and fifth AP pairs are plotted for WT (left) and Tg2576 mice (right) during the baseline or the presence of atropine. WT mice showed a significant increase in ISI between AP pairs during the baseline but Tg2576 mice did not. In the presence of atropine there the same was true, suggesting little effect of atropine.

When trains of 6 APs were analyzed (Figure 11C), there was adaptation in WT mice and atropine had no detectable effect, like the results from the 4 AP trains (two-way RMANOVA, F(1,143)=1.608, p=0.207; Figure 11D1). In Tg2576 mice, there was neither a significant effect of atropine (two-way RMANOVA, F(1,138)=0.534, p=0.466; Figure 11D1). When we compared the first and fifth AP pairs in WT mice, there was adaptation before as well as post-atropine (paired t-test, t=3.478, df=7, p=0.010; Figure 11D2), suggesting that atropine did not have an effect. In Tg2576 mice, there was no significant effect of atropine when the first and fifth AP pairs were analyzed (paired t-test, t=0.623, df=7, p=0.553; Figure 11D2). Thus, atropine had little effect on spike frequency adaptation. The small effects of atropine on adaptation in WT mice can be explained by a role of K^+^ currents other than M-current in spike frequency adaptation (e.g., I_A_, I_D_, I_C_; (Aiken et al., 1995; Storm, 1990). The lack of effect of atropine in Tg2576 mice is probably due to the very weak adaptation in the baseline conditions.

### III. GC morphology and location

#### A. Morphology

Confirmation that recorded cells were GCs was important because several types of GABAergic neurons are present in the GC layer (GCL; Hosp et al., 2014; Houser, 2007; Scharfman, 1995). Recorded cells were filled with biocytin by including it in the internal solution (see Methods). A randomly-selected group of recorded cells were evaluated by processing them to visualize biocytin (n=39; see Methods). An example of a filled GC is shown in Figure 12A3, and its morphology is consistent with a normal adult GC. GC somata were round or oval, and size (8-10 μm long) corresponded to past reports of rodent GCs (Amaral et al., 2007; Pierce et al., 2011; Rahimi and Claiborne, 2007). The dendrites extended throughout the molecular layer and there were numerous spines, similar to mature GCs (Amaral et al., 2007; Frotscher et al., 2000; Rahimi and Claiborne, 2007). The intrinsic properties also suggested that the recorded cells were GCs because intrinsic properties were consistent with previous studies of GCs rather than GABAergic neurons (Table 1; Scharfman, 1992).

**Figure 12.**
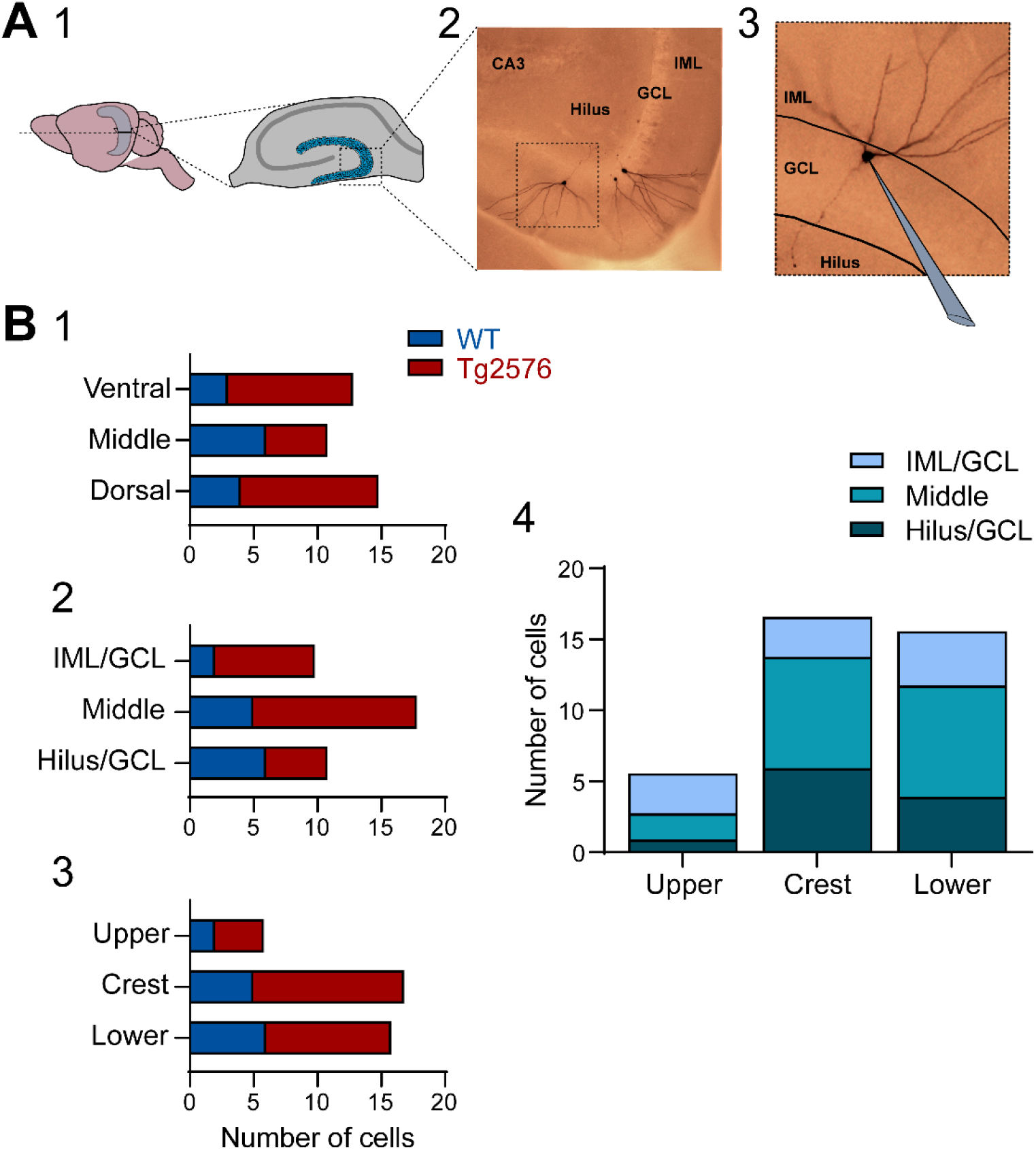
Morphology and location of sampled GCs in WT and Tg2576 mice. (A1) A diagram shows the plane that was used to generate horizontal slices. The diagram shows only one site in the septotemporal axis, but slices were selected from different parts of the septotemporal axis. (A2, A3) Examples of GCs labeled with biocytin are shown. The GC outlined by the box in A2 is expanded in A3. (B1-B3) The GCL was divided into a portion near the IML, a central portion and a portion close to the hilus. The differences between WT and Tg2576 GCs were not statistically significant. (B4) The data used in B3 are presented in an alternate way. Differences between WT and Tg2576 GCs were not statistically significant.

#### B. Position throughout the septotemporal axis

To determine if there was a preference for the WT GCs to be in a particular part of the DG that was distinct from Tg2576 mice, slices were examined to determine if they were from the dorsal, mid, or ventral DG. For the purposes of this comparison, the dorsal DG was defined as horizontal sections approximately 3.0 to 3.7 mm below the cortical surface, ventral DG as horizontal sections between approximately 4.3 to 5.0 mm, and the middle of the DG was defined as the area between the dorsal and ventral sites. As shown in Figure 12B1, there was no difference in the proportions of WT GCs recorded from dorsal, mid or ventral DG when they were compared to GCs from Tg2576 mice by Fisher’s exact test (all p>0.206).

#### C. Position within the GCL

We also compared the location of GCs in different subdivisions of the GCL (Figure 12B2). This was done to address whether GCs might be adult-born GCs, which are usually located near the hilus (Gould and Cameron, 1996), or semilunar GCs which are located near or in the IML (Williams et al., 2007). Therefore, we compared three subdivisions of the GCL: the GCL border with the IML, center of the GCL, and GCL border with the hilus (Figure 12B2). We found that similar proportions of GCs in WT and Tg2576 mice were located in these three parts of the GCL (Fisher’s exact test, all p>0.183; Figure 12B2).

#### D. Position within the DG blades

Next, we analyzed whether the GCs were preferentially localized to the upper blade, crest or lower blade, defining these areas as before (Bernstein et al., 2020). Figure 12B3 shows that the GCs from WT and Tg2576 mice did not appear to differ in the frequency they were recorded in the upper blade, crest, or lower blade. For statistical comparisons, WT was compared to Tg2576 for each location (Fisher’s exact test, all p>0.721).

### IV. Sex differences

Female and male mice were compared for most measurements described above. Regarding excitatory and inhibitory synaptic activity, WT males and females showed no statistical differences (Table 3). However, there were differences in male and female Tg2576 mice, where the females showed greater EPSC and nsC amplitudes than males (EPSCs: males: 5.25 ± 0.58 pA; females: 7.46 ± 0.54, unpaired t-test, t=2.761, df=11; p=0.019. NsC: males: 5.65 ± 0.57 pA; females: 7.69 ± 0.63, unpaired t-test, t=2.414, df=11; p=0.034; Table 3). intrinsic properties did not appear to differ (Table 4) but there were some notable differences in APs. The maximum rate of rise was significantly faster in females (males: 198.13 ± 23.05 mV/msec; females: 274.21 ± 23.47, unpaired t-test, t=2.160, df=17; p=0.045) and the same was true for the maximum rate of decay (males: −65.01 ± 4.30 mV/msec; females: −81.43 ± 2.76; unpaired t-test, t=2.710, df=17; p=0.015). However, the ratio of rise:decay did not change (males: 2.97 ± 0.21; females: 3.36 ± 0.27; unpaired t-test, t=1.156, df=17; p=0.264; Table 4).

**Table 3.** Sex differences in GCs synaptic properties. **Table 3 Legend.** Values of synaptic properties obtained from WT and Tg2576 GCs from male and female mice. (1) Unpaired student’s t-test for parametric data, and (2) Mann-Whitney test for non-parametric data.

| Synaptic properties<br>Males vs. Females |  |  |  |  |  |  |  |  |
| --- | --- | --- | --- | --- | --- | --- | --- | --- |
| Frequency | EPSPs |  | IPSCs |  | EPSCs |  | nsCs |  |
|  | WT | Tg2576 | WT | Tg2576 | WT | Tg2576 | WT | Tg2576 |
| Male | $1.56 \pm 0.12$ | $2.12 \pm 0.16$ | $10.38 \pm 1.01$ | $8.31 \pm 0.59$ | $3.16 \pm 0.33$ | $4.04 \pm 0.34$ | $0.10 \pm 0.04$ | $0.71 \pm 0.15$ |
| Female | $1.62 \pm 0.14$ | $2.54 \pm 0.46$ | - | $8.08 \pm 1.17$ | - | $4.10 \pm 0.50$ | - | $0.53 \pm 0.10$ |
| P value | 0.768 | 0.288 | - | 0.86 | - | 0.926 | - | 0.358 |
| Statistical Test | (1) | (1) | - | (1) | - | (1) | - | (1) |
| t, df | $t=0.301$, $df=14$ | $t=1.097$, $df=17$ | - | $t=0.181$, $df=12$ | - | $t=0.095$, $df=11$ | - | $t=0.960$, $df=11$ |
| N<sub>male</sub> | 9 | 13 | 13 | 7 | 7 | 7 | 12 | 7 |
| N<sub>female</sub> | 7 | 6 | - | 7 | - | 6 | - | 6 |

**Table 3 Legend.** Values of synaptic properties obtained from WT and Tg2576 GCs from male and female mice. (1) Unpaired student's t-test for parametric data, and (2) Mann-Whitney test for non-parametric data.
| Amplitude | EPSPs |  | IPSCs |  | EPSCs |  | nsCs |  |
| --- | --- | --- | --- | --- | --- | --- | --- | --- |
|  | WT | Tg2576 | WT | Tg2576 | WT | Tg2576 | WT | Tg2576 |
| Male | $0.48 \pm 0.04$ | $0.39 \pm 0.04$ | $10.90 \pm 1.11$ | $9.67 \pm 0.91$ | $8.72 \pm 1.03$ | $5.25 \pm 0.58$ | $4.91 \pm 0.79$ | $5.65 \pm 0.57$ |
| Female | $0.42 \pm 0.04$ | $0.42 \pm 0.05$ | - | $11.46 \pm 1.50$ | - | $7.46 \pm 0.54$ | - | $7.69 \pm 0.63$ |
| P value | 0.333 | 0.63 | - | 0.326 | - | **0.019** | - | **0.034** |
| Statistical Test | (1) | (1) | - | (1) | - | (1) | - | (1) |
| t, df | $t=1.002$, $df=14$ | $t=0.491$, $df=17$ | - | $t=1.024$, $df=12$ | - | $t=2.761$, $df=11$ | - | $t=2.414$, $df=11$ |
| N<sub>male</sub> | 9 | 13 | 13 | 7 | 7 | 7 | 12 | 7 |
| N<sub>female</sub> | 7 | 6 | - | 7 | - | 6 | - | 6 |

**Table 4.** Sex differences in GCs intrinsic properties. **Table 4 legend.** Values of the different intrinsic properties obtained from WT and Tg2576 GCs from male and female mice. Statistical comparisons were made of female WT vs. male WT and female Tg2576 vs. male Tg2576 GCs. (1) Unpaired student’s t-test for parametric data, and (2) Mann-Whitney test for non-parametric data.

|  |  | Intrinsic properties<br>Males vs. Females |  |  |  |  |  |  |  |
| --- | --- | --- | --- | --- | --- | --- | --- | --- | --- |
|  |  | RMP |  | R in (-5 pA) |  | Threshold |  | Peak Amplitude |  |
|  |  | WT | Tg2576 | WT | Tg2576 | WT | Tg2576 | WT | Tg2576 |
| Male |  | -69.00 ± 2.05 | -75.45 ± 2.94 | 350.64 ± 17.69 | 243.06 ± 27.74 | -37.92 ± 2.66 | -34.80 ± 1.97 | 104.46 ± 3.10 | 102.42 ± 4.27 |
| Female |  | -74.23 ± 3.01 | -79.55 ± 1.90 | 326.15 ± 32.11 | 228.25 ± 21.52 | -36.15 ± 2.14 | -40.21 ± 1.98 | 105.46 ± 3.09 | 112.00 ± 3.35 |
| P value |  | 0.165 | 0.312 | 0.539 | 0.788 | 0.670 | 0.089 | 0.842 | 0.096 |
| Statistical Test |  | (1) | (1) | (1) | (1) | (1) | (1) | (1) | (1) |
| t, df |  | t=1.456, df=16 | t=1.040, df=18 | t=0.633, df=12 | t=0.274, df=13 | t=0.434, df=16 | t=1.803, df=17 | t=0.203, df=16 | t=1.764, df=16.94 |
| N<sub>male</sub> |  | 12 | 12 | 8 | 11 | 12 | 12 | 12 | 12 |
| N<sub>female</sub> |  | 6 | 8 | 6 | 4 | 6 | 7 | 6 | 7 |

|  |  | Time to Peak |  | Time to Peak from threshold |  | Half-width |  | AHP |  |
| --- | --- | --- | --- | --- | --- | --- | --- | --- | --- |
|  |  | WT | Tg2576 | WT | Tg2576 | WT | Tg2576 | WT | Tg2576 |
| Male |  | 530.73 ± 96.62 | 576.17 ± 83.49 | 1.13 ± 0.12 | 1.17 ± 0.11 | 0.28 ± 0.02 | 0.32 ± 0.03 | -17.53 ± 1.51 | -16.38 ± 1.56 |
| Female |  | 449.70 ± 110.54 | 608.14 ± 91.81 | 0.88 ± 0.18 | 1.48 ± 0.07 | 0.26 ± 0.03 | 0.34 ± 0.02 | -16.34 ± 2.80 | -14.66 ± 1.68 |
| P value |  | 0.615 | 0.809 | 0.260 | 0.067 | 0.535 | 0.990 | 0.687 | 0.466 |
| Statistical Test |  | (1) | (1) | (1) | (1) | (1) | (2) | (1) | (1) |
| t, df |  | t=0.513, df=16 | t=0.246, df=17 | t=1.168, df=16 | t=1.959, df=17 | t=0.634, df=16 | Mann-Whitney U = 41 | t=0.411, df=16 | t=0.748, df=14.81 |
| N<sub>male</sub> |  | 12 | 12 | 12 | 12 | 12 | 12 | 12 | 12 |
| N<sub>female</sub> |  | 6 | 7 | 6 | 7 | 6 | 7 | 6 | 7 |

**Table 4 legend.** Values of the different intrinsic properties obtained from WT and Tg2576 GCs from male and female mice.
|  |  | Max Rise Slope |  | Max Decay Slope |  | dv/dt ratio |  | Tau |  |
| --- | --- | --- | --- | --- | --- | --- | --- | --- | --- |
|  |  | WT | Tg2576 | WT | Tg2576 | WT | Tg2576 | WT | Tg2576 |
| Male |  | 266.02 ± 28.01 | 198.13 ± 23.05 | -73.32 ± 4.69 | -65.01 ± 4.30 | 3.61 ± 0.22 | 2.97 ± 0.21 | 30.73 ± 2.80 | 21.72 ± 2.26 |
| Female |  | 256.24 ± 29.78 | 274.21 ± 23.47 | -75.28 ± 5.26 | -81.43 ± 2.76 | 3.37 ± 0.28 | 3.36 ± 0.27 | 26.06 ± 1.99 | 21.02 ± 1.28 |
| P value |  | 0.831 | **0.045** | 0.703 | **0.015** | 0.517 | 0.264 | 0.288 | 0.828 |
| Statistical Test |  | (1) | (1) | (1) | (1) | (1) | (1) | (1) | (1) |
| t, df |  | t=0.217, df=16 | t=2.160, df=17 | t=0.388, df=16 | t=2.710, df=17 | t=0.662, df=16 | t=1.156, df=17 | t=1.099, df=16 | t=0.221, df=17 |
| N<sub>male</sub> |  | 12 | 12 | 12 | 12 | 12 | 12 | 12 | 12 |
| N<sub>female</sub> |  | 6 | 7 | 6 | 7 | 6 | 7 | 6 | 7 |

## DISCUSSION

### I. Synaptic activity in WT and Tg2576 mice

The results from our studies of synaptic activity suggest that sEPSPs and sEPSCs were greater and sIPSPs were reduced in Tg2576 mice compared to WT. For sEPSPs and sEPSCs, the frequency increased, particularly of small events. The mean amplitude decreased, although it was significant only for sEPSCs. For sIPSPs, frequency was reduced, not amplitude. Together these changes suggest a potential reason why the Tg2576 mouse was found previously to exhibit increased excitability at early ages (Bezzina et al., 2015; Kam et al., 2016): GCs showed increased excitation and decreased inhibition. Having said that, it is recognized that the DG might be only one of several areas contributing to increased excitability in hAPP mice (Bezzina et al., 2015; Kam et al., 2016; Palop et al., 2007; Verret et al., 2012; You et al., 2017). However, a focus on the DG is appropriate given the DG has been implicated in the pathophysiology of mouse models of AD neuropathology (see Introduction). Moreover, DG and CA3 hyperactivity have been implicated in early stages of AD (Bakker et al., 2012). In addition, the DG is considered to be a critical regulator of hippocampal hyperexcitability in epilepsy research (Heinemann et al., 1992; Krook-Magnuson et al., 2015; Lothman et al., 1992; Scharfman, 2019). Notably, our data are consistent with the enhanced expression of markers of neuronal activity observed in GCs in another mouse model of human APP mutations (J20 mice) (You et al., 2017).

There could be many reasons why sEPSP and sEPSC frequency was high in Tg2576 GCs relative to WT GCs. One possibility is that there was increased activity in glutamatergic afferents to the GCs. An attractive candidate is the perforant path input from the entorhinal cortex because it is considered to be the major excitatory input to the DG (Amaral et al., 2007). Also, the entorhinal cortex is highly vulnerable in AD, exhibiting early pathology (Braak and Braak, 1991; Kordower et al., 2001) especially in the perforant path connections to the DG (de Leon et al., 2007; de Toledo-Morrell et al., 2000). In the Tg2576 mouse, we found that there also are early signs of degenerative changes, and there was increased excitability in slices of the entorhinal cortex at 2-4 months of age (Duffy et al., 2015). Another study also found increased excitability at early ages in Tg2576 mice using *in vivo* recordings of entorhinal cortical units (Xu et al., 2015). It would be logical that increased entorhinal cortical excitability could contribute to an increase in perforant path activity, leading to increased excitatory input to the GC dendrites. This would potentially contribute to the increased frequency of sEPSPs and sEPSCs observed in the present study.

Notably, a previous study showed that at 4 months of age, GCs of Tg2576 mice exhibited reduced spines (Jacobsen et al., 2006). This result might be a response to overactivity at early ages because spine loss is observed in GCs in response to hyperexcitability (Isokawa, 2000) and in cultures after increased excitation due to GABA receptor antagonism (Drakew et al., 1996). Interestingly, long-term potentiation of the perforant path was impaired at 5 months of age in the Tg2576 mice with reduced spines (Jacobsen et al., 2006), consistent with a reduction in spines in other animals when LTP is impaired (D’Agostino et al., 2013). These studies are important to consider together because they suggest that early hyperexcitability precedes and possibly contributes to impairments later in life which has been hypothesized before (Kam et al., 2016).

There are other possible contributing factors that explain why sEPSPs and sEPSCs increased in GCs in Tg2576 mice. For example, a previously undetectable input might become detectable. This could apply to the perforant path, but it also could be another input. It seems unlikely that altered intrinsic properties were the cause, because R_in_ decreased in Tg2576 mice, and decreased R_in_ usually reduces the sensitivity of a neuron to its afferent input. Instead, one possible cause of increased sensitivity is that the normal role of muscarinic receptors to suppress glutamate release was reduced in Tg2576 mice. If transmitter release increased, frequency would rise. This role of muscarinic receptors could also help explaining why there were increased nsCs in Tg2576 mice relative to WT mice (discussed further below). This idea is attractive because it explains how changes in cholinergic input can alter the DG in Tg2576 mice and possibly other hAPP mice.

This possible role of muscarinic receptors in regulating excitatory input to GCs is consistent with prior studies. Septohippocampal neurons are the major cholinergic input to the DG and hippocampus (Aznavour et al., 2005; Clarke, 1985; Deller et al., 1999; Dougherty and Milner, 1999; Frotscher, 1991; Frotscher and Leranth, 1985; Frotscher and Leranth, 1986; Leranth and Frotscher, 1987; Milner and Veznedaroglu, 1993; Nyakas et al., 1987; Takacs et al., 2018; Wainer et al., 1985) and normally one of the roles of acetylcholine is to suppress release of transmitter from external inputs to the DG and hippocampus, a suppression that occurs during sleep and allows more emphasis on circuits intrinsic to the DG/hippocampus and memory consolidation (Hasselmo et al., 1996). However, in the Tg2576 mouse there appears to be early abnormalities of the septohippocampal input (Kam et al., 2016). Specifically, septal neurons appear to be overactive in that they express c-Fos (Kam et al., 2016). In addition, young Tg2576 mice have increased hippocampal expression of the synthetic enzyme for acetylcholine, choline acetyl transferase (ChAT), which is present in cholinergic fibers from cholinergic neurons extrinsic to the DG (Kam et al., 2016). ChAT also rises early in AD (DeKosky et al., 2002; Ikonomovic et al., 2007). One possibility is that muscarinic receptors were downregulated in young Tg2576 mice in response to increased release of acetylcholine from septohippocampal nerve terminals. As a result, Tg2576 mice would have less cholinergic suppression of glutamate release.

The finding in the present study that sIPSCs were reduced in Tg2576 mice could also contribute to the increased frequency of sEPSPs and sEPSCs. Reduced sIPSCs is consistent with prior studies which suggest that the GABAergic neurons in mice with human APP mutations develop deficient sodium channels (Nav1.1) which make them less able to release GABA (Martinez-Losa et al., 2018; Verret et al., 2012). Another possibility is that the deficient sodium channels occur later in life, and the first defect in sIPSCs is due to another mechanism. One early mechanism would be alteration of the normal effects of septohippocampal terminals on GABAergic neurons. Normally acetylcholine has diverse effects on GABAergic neurons which depend in part on behavioral state and in part on the type of GABAergic neurons (Behrends and ten Bruggencate, 1993; Chiang et al., 2010; Dannenberg et al., 2017; Dougherty and Milner, 1999; Frazier et al., 1998; Frazier et al., 2003; Leranth and Frotscher, 1987; McQuiston and Madison, 1999; Pabst et al., 2016; Pitler and Alger, 1992; Raza et al., 2017; Yi et al., 2014), making a clear mechanism hard to propose at the present time.

### II. Intrinsic properties and firing behavior of WT and Tg2576 mice

The results showed that intrinsic properties and firing behavior were altered in Tg2576 mice relative to WT. RMP was more hyperpolarized, and both R_in_ and τ were reduced in Tg2576 mice. Also, APs in Tg2576 GCs had a longer time to peak and a longer half-width. firing behavior showed that adaptation in Tg2576 GCs was weak. Together these differences suggest complex changes in GCs in young Tg2576 mice. While weaker adaptation would make a cell more likely to fire APs, a more hyperpolarized RMP and reduced R_in_ would potentially decrease the ability of afferent input to influence the cell. The predicted net effect would be to reduce the ability of GCs to respond to afferent input while increasing the ability of GCs to fire APs.

If this is true, it would be likely to cause substantial network defects because APs would fire outside of the times they would normally be triggered by afferent input. The DG would become unlikely to play its normal role in cognition and behavior because it would fire at inappropriate times. In addition, the longer time to peak and half-width of Tg2576 GC APs might also render GCs less effective in their normal roles because APs would be slower to peak. The slow time-to-peak could lead to delays in the timing of AP discharge and transmitter release.

Together the results from synaptic and intrinsic studies of GCs suggest possible cellular mechanisms for decreased cognitive function in Tg2576 mice. The idea that cognitive impairments develop early in life (2-4 months-old) is supported by a prior study showing worse performance of approximately 3 month-old Tg2576 mice on tasks that test spatial memory (e.g. novel object location; Duffy et al. 2015). In the CRND8 mouse, which has two human APP mutations (APP_Swe_, APP_Ind_), the novel object location task has also been found to be deficient, and the animals were just2 months of age (Francis et al., 2012). In the J20 mouse, the novel object location task is impaired early in life too (You et al., 2017). Notably, task performance was improved by specific experimental manipulations of GCs (You et al., 2017), implicating the DG in cognitive impairments.

It is remarkable that the changes in synaptic events and intrinsic properties in Tg2576 mice occurred so early in life. It begs the question how Aβ could trigger effects so early and how there could be so many effects. One potential answer is that effects of Aβ are not the only cause, but APP and APP metabolites also contribute. This idea has support from an earlier study of Tg2576 mice (Xu et al., 2015) as well as a wealth of evidence that APP metabolites other than Aβ cause numerous adverse effects (Nixon, 2017). Regarding the diversity of effects, this might be due to the fact that some effects are primary whereas others are compensatory. For example, there may first be an increase in excitability due to greater activity of excitatory input with reduced inhibitory input, and then a hyperpolarization and reduction in R_in_ to compensate.

### III. Atropine in WT and Tg2576 mice

#### A. Synaptic activity

##### 1) sEPSPs, sEPSCs and sIPSCs

Atropine had diverse effects in Tg2576 mice compared to WT mice, sometimes being similar and sometimes quite different. WT and Tg2576 mice were similar in the effect of atropine to reduce sIPSC frequency and amplitude. WT and Tg2576 mice were different in the effects of atropine on excitatory events. For sEPSPs, WT mice showed an increase in frequency in response to atropine, but this was not evident in Tg2576 mice. One reason, already mentioned in the Results, could be that Tg2576 mice had a higher baseline frequency of sEPSPs than WT mice, and atropine could not increase sEPSP frequency further. This explanation would lead to the implication that altered muscarinic receptors were in part responsible for the high baseline frequency of sEPSPs in Tg2576 mice. What alteration in muscarinic receptors might occur is not clear. An attractive mechanism would be that muscarinic receptors are slightly impaired or downregulated by hyperactivity of cholinergic inputs early in life. Thus, high concentrations of acetylcholine could cause a compensatory internalization of muscarinic receptors. Hyperactivity of septal cholinergic inputs in young Tg2576 mice is supported by a prior study showing that there was high c-Fos protein expression in medial septal neurons in young Tg2576 mice (Kam et al., 2016).

##### 2) nsCs

Atropine also had a very interesting effect in WT and Tg2576 mice when GC holding potential was 0 mV. Small inward currents were present, especially in Tg2576 mice. Atropine increased these events in WT mice and had a small effect in Tg2576 mice. These events, which we call nsCs, were glutamatergic based on blockade by ionotropic receptor antagonists. NsCs may have contributed to the increase in sEPSP and sEPSC frequency (Figures 1, 2) but were simply much clearer using voltage clamp at 0 mV. They are potentially important because they suggest another way the cholinergic system is abnormal in young Tg2576 mice and could contribute to increased excitability of GCs.

The mechanisms underlying nsCs and the effect of atropine could be pre- or postsynaptic. One possible mechanism is that some excitatory afferents that normally are a small excitatory input become enhanced, particularly in Tg2576 mice. Furthermore, these inputs were increased by atropine, suggesting that normally muscarinic receptors led to a tonic suppression of nsCs. There are two likely candidate inputs because GCs primarily have two glutamatergic inputs, the perforant path and mossy cells. Regarding mossy cells, recent data suggest they have a small excitatory effect on GCs (Bernstein et al., 2020). They are hard to consider for a potential role in nsCs because muscarinic receptors are present on mossy cells but how they regulate the terminals of mossy cells is not clear (Hofmann and Frazier, 2010).

It seems likely that the perforant path would contribute to nsCs because it is known that the entorhinal cortex exhibits hyperexcitability in young Tg2576 mice (Duffy et al., 2015; Xu et al., 2015). We showed increased excitability in slices, and also showed early changes in neuronal architecture (Duffy et al., 2015). Xu et al. (2015) found evidence of hyperexcitability in the lateral entorhinal cortex using unit recording *in vivo*. If hyperexcitability in the entorhinal cortex leads to more activity in the perforant path, the result would be increased excitatory input to GCs. Importantly, the entorhinal cortex receives extensive innervation by septal cholinergic afferents, making it more likely that the perforant path was responsible for greater inward currents in Tg2576 mice and a release from suppression in WT mice by atropine. These findings support the view that the cholinergic system in Tg2576 mice is altered at very early ages in multiple ways.

##### 3) Intrinsic properties and firing behavior

Atropine also had effects on intrinsic properties that differed between WT and Tg2576 mice. Thus, atropine reduced R_in_ but only in Tg2576 mice. Atropine made threshold more depolarized in Tg2576 mice, but did not in WT. The difference between WT and Tg2576 threshold would make the Tg2576 GCs less able to fire APs. Together these effects help explain the ability of atropine to decrease aberrant interictal spikes *in vivo* in the Tg2576 mice (Kam et al., 2016). On the other hand, if atropine induced nsCs and decreased sIPSCs, atropine would be likely to increase excitability.

Thus, a remaining question is how the results might explain the effect of atropine to decrease IIS *in vivo* (Kam et al., 2016). In the GCs, atropine increased excitability by increasing nsCs, and by depressing IPSCs. It also had effects on intrinsic properties and firing behavior that might have decrease excitability. One reason is that many of the animals tested with atropine in the *in vivo* study by Kam et al. (2016). were older than 2-3 months. At that time, atropine might have less effect on afferent input because of the gradual reduction in entorhinal, cholinergic and GABAergic function caused by Aβ pathology. That would leave the effects of atropine on intrinsic properties and those effects reduced excitability.

This discussion is potentially relevant to other cholinergic treatment in AD. Thus, individuals with AD that are treated with drugs that act on cholinergic synapses and receptors, such as donepezil, may exert different effects depending on the age and stage of pathology. It could explain the clinical data showing that some individuals are helped by these drugs, and others are not (Hampel et al., 2018; Richter et al., 2018).

### IV. Sex differences

Sex differences were useful to clarify because of widespread observations that female mice with hAPP mutations are often more affected than males (Bangasser et al., 2017; Clinton et al., 2007; Jiao et al., 2016; Roy et al., 2018; Yang et al., 2018). In the present study, females had larger sEPSCs and larger nsCs in Tg2576 mice. Females also had greater maximum rate of rise and rate of decay of APs in Tg2576 mice. These effects might make females more prone to excitation, leading to hyperexcitability. The hyperexcitability would be consistent with a worse phenotype in females than males in prior studies. A reason why hyperexcitability would be associated with a worse phenotype is that hyperexcitability can lead to overexcitation of downstream targets. Overexcitation of the DG and CA3 may cause impaired cognition (Bakker et al., 2012). In addition, hyperexcitability of GCs can lead to excitotoxicity to hilar cells and CA3 pyramidal cells because the GC giant boutons can release high concentrations of glutamate (Acsady et al., 1998; Chicurel and Harris, 1992; Rama et al., 2019; Rollenhagen and Lubke, 2010; Scharfman, 2016; Scharfman and Bernstein, 2015). Over time hilar and CA3 cells might die, but if they did not die, they would be likely to become impaired by the metabolic demand to support the maintenance of normal function in the face of repeated exposure to high concentrations of glutamate (Choi, 1994; Olney et al., 1986; Sattler and Tymianski, 2001; Schwarcz et al., 1984).

### V. Role of Aβ

Aβ in its oligomeric (“soluble”) form may have contributed to the changes reported in this study. In addition, the precursor APP or APP metabolites could have contributed. APP could play a role because it is overexpressed from birth. However, to date none of the synaptic changes or intrinsic properties we identified have been shown to occur in response to APP. It is not clear when Aβ levels begin to rise in the Tg2576 mouse. However, Aβ levels were high by ELISA at just 3 months of age in Tg2576 mice (Duffy et al., 2015). Therefore, the results in this study could have been related to the earliest escalation in Aβ.

In the literature, there are diverse effects of adding APP, its metabolites, or Aβ to hippocampal slices. Although studies of intrinsic properties are rare, multiple studies have documented synaptic effects on hippocampal neurons and a few on GCs (Chen, 2005; Chen et al., 2002; Ciccone et al., 2019; Eslamizade et al., 2015; Hou et al., 2009; Pena et al., 2010; Rovira et al., 2002; Tamagnini et al., 2015; Wu et al., 1995). One of the problems in relating these papers to the results presented here is that it is hard to equate superfusion of synthetic peptides to endogenous production of Aβ from mutated APP. Nevertheless, the published data using exogenous application of Aβ peptides are consistent with what we have shown, because diverse changes in synaptic activity have been documented, both increases and decreases (Gulisano et al., 2019; Kelly et al., 1996; Parameshwaran et al., 2007; Ripoli et al., 2014; Yao et al., 2013).

### VI. Conclusions

This study provided insight into the early changes in GCs of the Tg2576 mouse model of Alzheimer’s neuropathology. These data help address the hypothesis that the DG is an area that is affected at early ages and participates in the progressive pathophysiology of the mouse model. By examining ages that had not been examined previously, and by using whole cell recording, detailed insight into synaptic and intrinsic properties were obtained. The results suggest diverse synaptic changes develop in GCs that could contribute to increased excitability and cognitive impairment. By showing intrinsic properties are affected the results are the first to show that intrinsic characteristics are important to study. Finally, by elaborating the effects of atropine the results support the hypothesis that the cholinergic system is altered in diverse ways at early ages. While most studies begin at ages much older than what we chose, the present data set is a contrast and supports the view that Alzheimer’s pathology begins long before the presence of amyloid plaques.

---

**Supplementary Figure 1.**
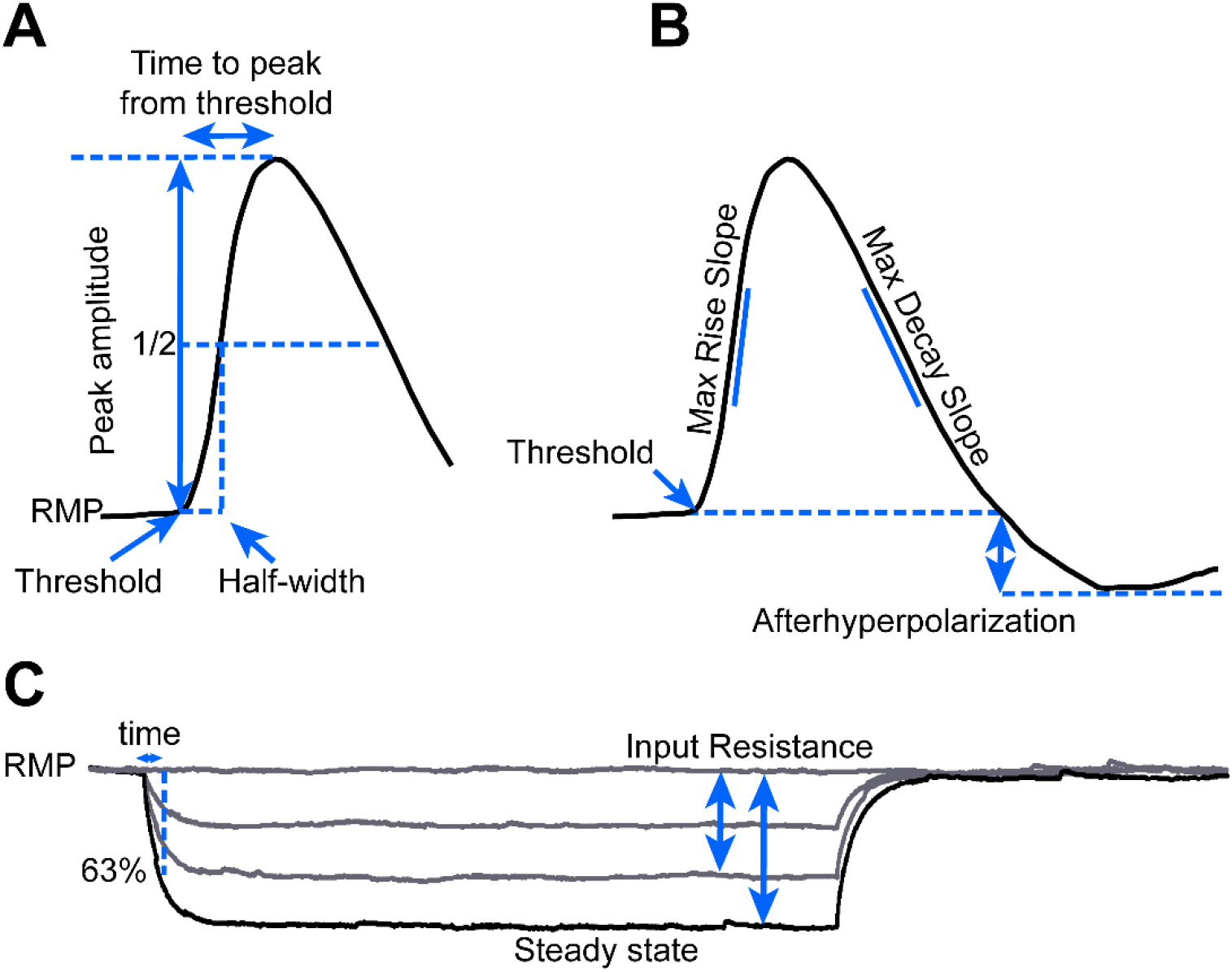
Measurement of intrinsic properties. (A) An example of an AP shows the measurement of AP threshold, half-width, peak amplitude, and time to peak from threshold. For these measurements an AP was evoked using a depolarizing current step. The current used for the step was the minimum that elicited an AP in approximately half the trials. (B) Measurements are shown for the maximum rise and decay slopes of the AP, and the afterhyperpolarization. (C) Representative traces are used to show the measurement of responses to current pulses used to determine R_in_. In addition, the measurements that were used to determine tau are shown. Tau was defined as the time when the response reached 63% of its steady state amplitude.

**Supplementary Figure 2.**
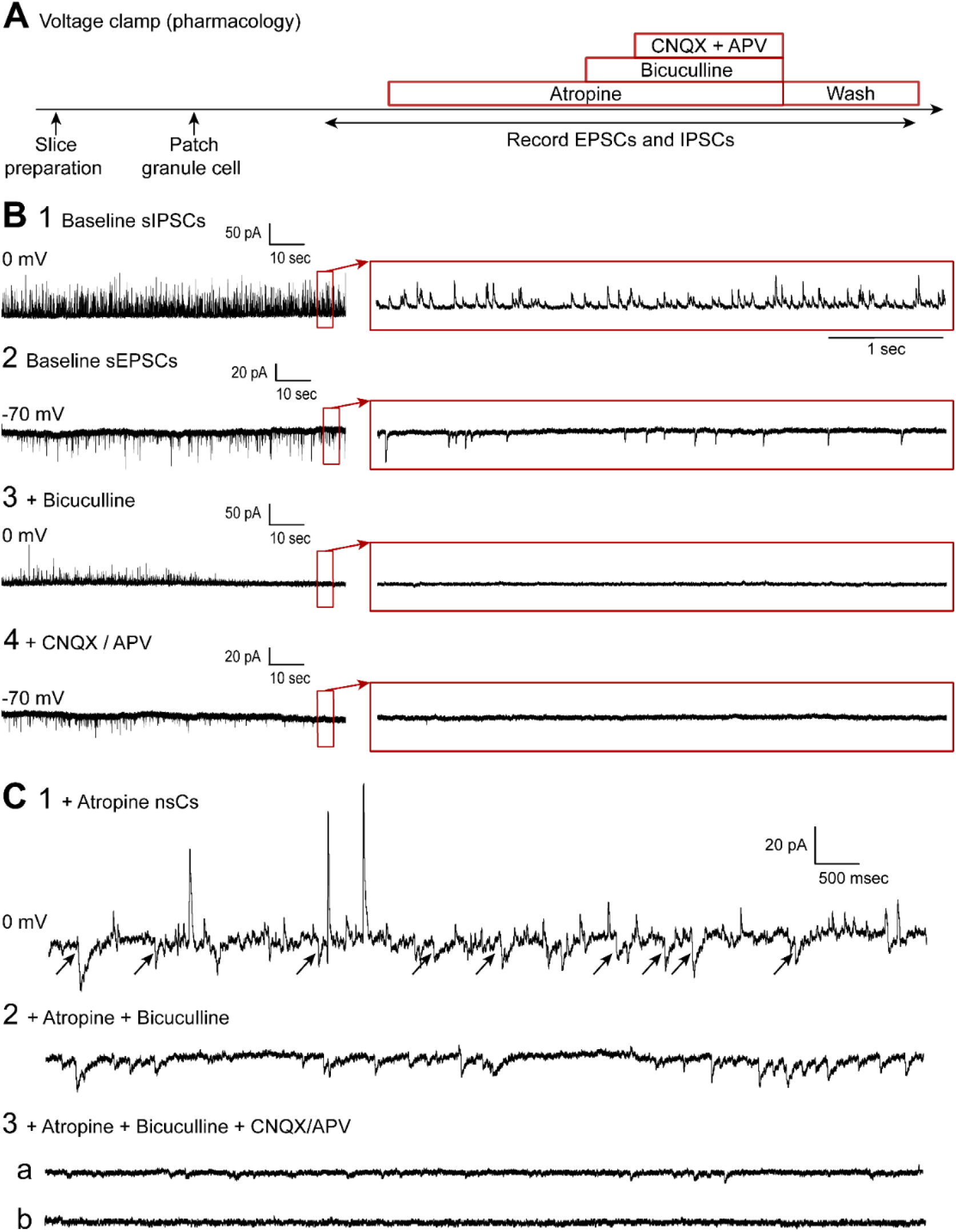
Confirmation that IPSCs were GABAergic and EPSCs and nsCs were glutamatergic. (A) Timeline for the pharmacological experiments to address the GABAergic or glutamatergic nature of synaptic activity. These recordings were conducted in the presence of atropine so that nsCs could be easily detected. Initially recordings in voltage clamp were conducted to establish a baseline. Then atropine (10 μM) was added, followed by bicuculline (10 μM), and subsequently CNQX (10 μM) and APV (50 μM). (B1) Representative traces of sIPSCs during the baseline. (B2) Representative traces of sEPSCs during the baseline. (B3) Bicuculline blocked sIPSCs. (B4) CNQX and APV blocked sEPSCs. Synaptic activity returned after a period of recording in drug-free ACSF (not shown). (C1) Examples of nsCs (arrows) in a Tg2576 GC recorded in the presence of atropine at holding potential of 0 mV. (C2) Bicuculline did not block nsCs, suggesting they were not mediated by GABAA receptors. (C3) CNQX and APV blocked nsCs, suggesting they were mediated by glutamate acting at ionotropic receptors.

## CREDIT STATMENTS

**David Alcantara-Gonzalez:** conceptualization, methodology, formal analysis, investigation, writing-original draft preparation/creation, visualization, project administration.

**Elissavet Chartampila:** investigation.

**Helen E. Scharfman:** conceptualization, methodology, resources, writing-review & editing, visualization, supervision, project administration, funding acquisition.

## ACKNOWLEDGEMENTS

We thank Dr. Aine Duffy for her contributions to the initiation of this project. We also thank Drs. Justin Botterill, Chiara Criscuolo, Christos Lisgaras and Yi-Ling Lu for discussion during the preparation of this paper. This project was supported by NIH R01 AG-055328 to H.E.S. and the New York State Office of Health.

## Notes

### Competing Interest Statement

The authors have declared no competing interest.

